# PUMILIO hyperactivity drives premature aging of *Norad*-deficient mice

**DOI:** 10.1101/432112

**Authors:** Florian Kopp, Mehmet E. Yalvac, Beibei Chen, He Zhang, Sungyul Lee, Frank A. Gillett, Mahmoud M. Elguindy, Sushama Sivakumar, Hongtao Yu, Yang Xie, Prashant Mishra, Zarife Sahenk, Joshua T. Mendell

## Abstract

Although numerous long noncoding RNAs (lncRNAs) have been identified, our understanding of their roles in mammalian physiology remains limited. Here we investigated the physiologic function of the conserved lncRNA *Norad in vivo.* Deletion of *Norad* in mice results in genomic instability and mitochondrial dysfunction, leading to a dramatic multi-system degenerative phenotype resembling premature aging. Loss of tissue homeostasis in Norad-deficient animals is attributable to augmented activity of PUMILIO proteins, which act as post-transcriptional repressors of target mRNAs to which they bind. *Norad* is the preferred RNA target of PUMILIO2 (PUM2) in mouse tissues and, upon loss of *Norad,* PUM2 hyperactively represses key genes required for mitosis and mitochondrial function. Remarkably, enforced *Pum2* expression fully phenocopies *Norad* deletion, resulting in rapid-onset aging-associated phenotypes. These findings provide new insights and open new lines of investigation into the roles of noncoding RNAs and RNA binding proteins in normal physiology and aging.

## INTRODUCTION

Long noncoding RNAs (lncRNAs) comprise a heterogeneous class of transcripts that are defined by a sequence length greater than 200 nucleotides and the lack of a translated open reading frame (ORF). lncRNAs have been proposed to perform a variety of cellular functions including regulation of gene expression in *cis* and *trans,* modulation of functions of RNAs and proteins to which they bind, and organization of nuclear architecture^1^. Although they have been estimated to number in the tens of thousands^2^, the biological significance of the vast majority of lncRNAs remains to be established. This is due, in part, to the generally low abundance and poor evolutionary conservation of most lncRNAs, which has limited our ability to interrogate their biochemical functions as well as their biologic roles *in vivo* using model organisms. Moreover, while genetic studies in mice have uncovered important functions for selected mammalian lncRNA loci in development and disease states^3,4^, it has often been challenging to connect specific lncRNA-driven phenotypes to defined RNA-mediated functions. As a result, our broad understanding of how the molecular pathways controlled by lncRNAs impact development and physiology remains limited.

*Noncoding RNA activated by DNA damage* (*NORAD*) is a recently described lncRNA that is distinguished from the majority of transcripts in this class due to its high abundance in mammalian cells and strong evolutionary conservation across mammalian species^5,6^. Studies in human cells have established that this RNA functions as a strong negative regulator of PUMILIO1 (PUM1) and PUMILIO2 (PUM2), RNA binding proteins (RBPs) that belong to the deeply conserved family of Pumilio and Fem3 binding factor (PUF) proteins. PUM1/2 bind specifically to the 8 nucleotide (nt) PUMILIO response element (PRE) (UGUANAUA), which is often located in the 3’ UTR of mRNAs. Binding of PUM1/2 to these sites triggers accelerated deadenylation, reduced translation, and turnover of mRNA targets^7,8^. With the capacity to bind a large fraction of PUM1/2 within the cell, *NORAD* limits the availability of these proteins to repress target mRNAs^5,6^. Consequently, inactivation of *NORAD* results in PUMILIO hyperactivity with augmented repression of a program of target mRNAs that includes key regulators of mitosis, DNA repair, and DNA replication. Dysregulation of these genes results in dramatic genomic instability in NORAD-deficient cells^5^. In accordance with this model, PUM2 overexpression is sufficient to phenocopy, while PUM1/2 loss-of-function is sufficient to suppress, the *NORAD* knockout phenotype in human cells. Recent work has also identified an interaction between *NORAD* and RBMX^9^, an RNA binding protein that contributes to the DNA damage response, but the role of this interaction in the control of genomic stability by *NORAD* remains unknown.

Although PUF proteins are deeply conserved across eukaryotic species, the emergence of *NORAD* specifically within mammals suggests the existence of strong selective pressure to maintain tight control of PUMILIO activity within this lineage. Interestingly, mammalian neurons are exquisitely sensitive to reduced dosage of PUMILIO, with only a 25% to 50% reduction in PUM1 expression resulting in neurodegeneration in both human and mouse brain^10,11^. The existence of *NORAD* suggests that hyperactivity of PUMILIO, expected to occur in the absence of this lncRNA, may also result in deleterious effects. While studies in cell lines have demonstrated that *NORAD* loss or PUMILIO overexpression results in genomic instability, the consequences of mammalian PUMILIO hyperactivity *in vivo* have yet to be examined.

Here, we report an investigation of the physiologic role of the Norad-PUMILIO axis through the generation and characterization of Norad-deficient and *Pum2* transgenic mouse lines. While deletion of *Norad* does not overtly impact development, mice lacking this lncRNA develop a dramatic multi-system degenerative phenotype that resembles premature aging. Loss of *Norad* results in PUMILIO hyperactivity and repression of genes that are essential for normal mitosis, leading to the accumulation of aneuploid cells in Norad-deficient tissues. Unexpectedly, *Norad* loss also causes striking mitochondrial dysfunction, associated with repression of PUMILIO targets that regulate mitochondrial homeostasis. Importantly, transgenic expression of *Pum2* is sufficient to fully phenocopy *Norad* loss of function, triggering genomic instability, mitochondrial dysfunction, and rapidly-advancing aging-associated phenotypes. These findings demonstrate the importance of *Norad* in maintaining tight control of PUMILIO activity *in vivo* and establish a critical role for this lncRNA-RBP regulatory interaction in mammalian physiology.

## RESULTS

### Deletion of the mouse *Norad* ortholog

A mouse ortholog of *NORAD* (*2900097C17Rik* or *Norad*), exhibiting 61% nucleotide identity with its human counterpart, is clearly identifiable on mouse chromosome 2 (Fig. 1a). Like the human transcript, mouse *Norad* shows minimal protein-coding potential as assessed by PhyloCSF, a metric that discriminates between coding and noncoding sequences based on their evolutionary signatures^12^ (Supplementary Fig. 1a). Both human and mouse *NORAD*/*Norad* are ubiquitously expressed throughout the body, with highest abundance in brain (Fig. 1b and Supplementary Fig. 1b), and are predominantly localized to the cytoplasm, as determined by cell fractionation studies in mouse embryonic fibroblasts (MEFs) (Fig. 1c) and human cell lines^5,6^.

**Fig. 1.**
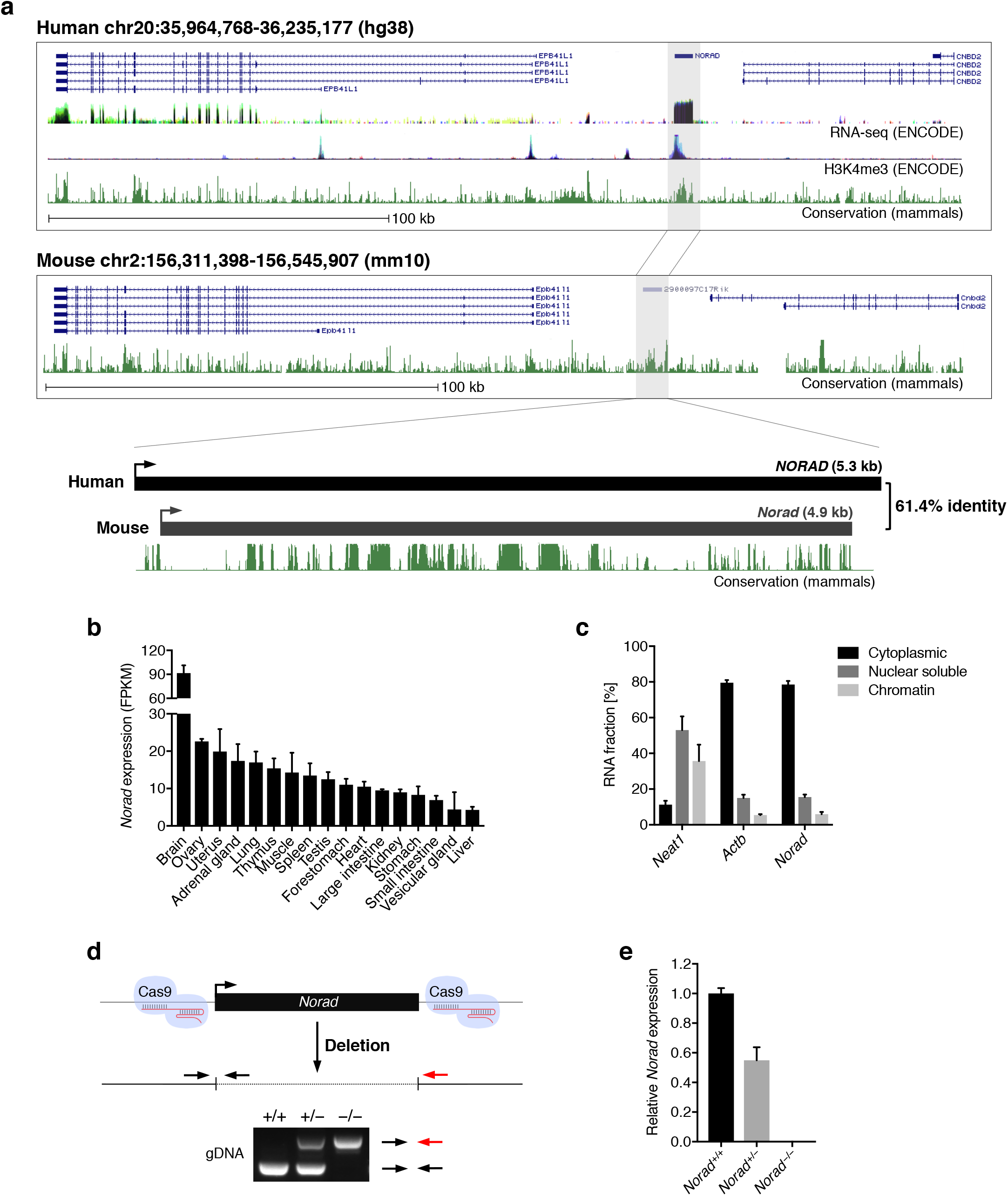
Deletion of the mouse *Norad* ortholog. **a**, Syntenic regions of human and mouse chromosomes 20 and 2, respectively, harboring *Norad* orthologs. ENCODE RNA-seq and H3K4me3 ChIP-seq coverage in human as well as conservation in mammals (PhastCons) are depicted as UCSC Genome Browser tracks. **b**, *Norad* expression across 17 mouse tissues based on RNA-seq data from the Mouse Transcriptomic BodyMap^48^. **c**, Predominantly cytoplasmic localization of *Norad* in immortalized MEFs. Subcellular localization was determined by cellular fractionation with subsequent qRT-PCR of *Norad*, *Neat1* (nuclear control), or *Actb* (cytoplasmic control) (n = 3 biological replicates). **d**, Schematic of CRISPR/Cas9-mediated genome-editing strategy used to delete *Norad* in mice. Genotyping PCR strategy and a representative genotyping result are shown. **e**, *Norad* expression in primary MEF lines of the indicated genotypes, determined by qRT-PCR (n = 3 biological replicates). Data are represented as mean ± SD in (**b**), (**c**), and (**e**).

To investigate the function of the mouse *Norad* ortholog, clustered regularly interspaced short palindromic repeats (CRISPR)/Cas9-mediated genome editing was employed to delete the lncRNA-encoding sequence from the mouse genome, yielding three independent knockout lines (Fig. 1d, e and Supplementary Fig. 1c). Importantly, expression of the neighboring genes *Epb41l1* and *Cnbd2* was unaffected in *Norad^−/−^* brain and spleen (Supplementary Fig. 1d), demonstrating that deletion of the *Norad* locus does not perturb local gene regulation.

### *Norad* loss-of-function results in a degenerative phenotype resembling premature aging

*Norad^−/−^* mice were viable, fertile, and born at the expected Mendelian frequency (Supplementary Fig. 2a). Early in life, *Norad*-deficient mice were indistinguishable from wild-type littermates, but by six months of age, the onset of a multi-system degenerative phenotype resembling premature aging became apparent, with approximately 50% of the mice developing severe manifestations by one year of age (Fig. 2a, b). This phenotype was characterized by accelerated alopecia and graying of fur in male *Norad^−/−^* mice (Fig. 2a and Supplementary Fig. 2b), while both male and female *Norad^−/−^* mice displayed pronounced kyphosis (Fig. 2b and Supplementary Fig. 2c). Increased kyphosis was also evident in *Norad^+/–^* animals, indicating a dose-dependent effect of *Norad* loss of function. Although body weight was comparable between cohorts of *Norad^+/+^*, *Norad^+/–^*, and *Norad^−/−^* mice at one year of age (Supplementary Fig. 2d), mice that developed outward features of aging such as kyphosis also exhibited significant weight loss accompanied by loss of total body fat and subcutaneous fat (**Fig. 2c-e**). Abnormalities were also apparent in *Norad*-deficient skeletal muscle, with marked switching of fast twitch glycolytic (FTG) fibers to slow twitch oxidative (STO) fibers (**Fig. 2f**), a phenomenon associated with normal muscle aging^13^. In addition, *Norad^−/−^* mice showed accelerated onset of aging-associated pathologies within the central nervous system (CNS)^14–16^, including condensed neuronal cell bodies and an accumulation of lipofuscin and vacuoles within spinal motor neurons (Fig. 2g and Supplementary Fig. 2e). Similar pathologies were evident in neurons in the dorsal root ganglia, brain stem, and cerebellum (data not shown). These changes were accompanied by an overall reduction in neuronal density in the spinal cord (Supplementary Fig. 2f). Finally, *Norad^−/−^* mice showed decreased overall survival over a two-year interval (Fig. 2h). These findings demonstrate that *Norad* is essential to suppress widespread aging-associated degenerative phenotypes in mice.

**Fig. 2.**
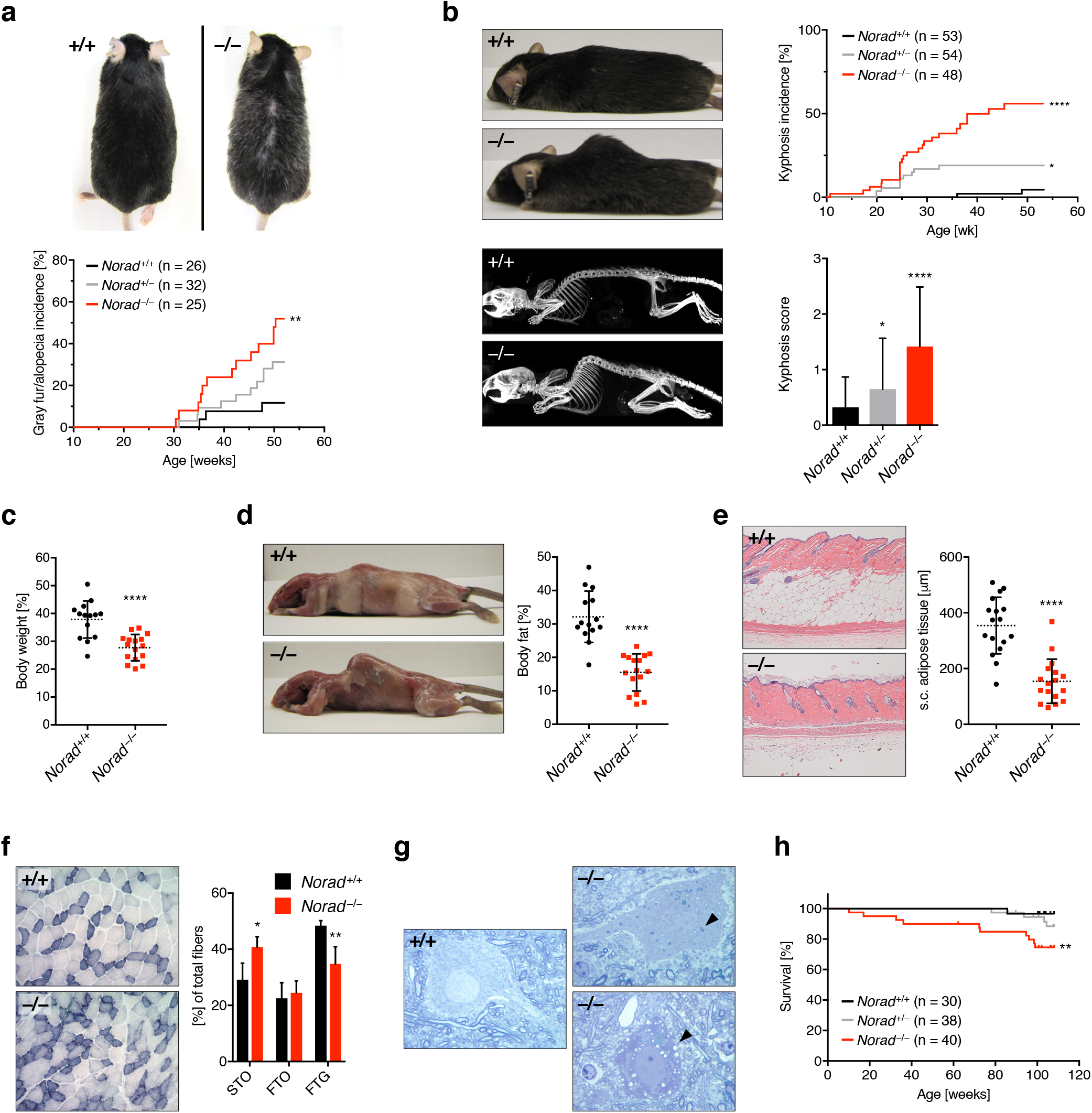
*Norad* loss-of-function results in a degenerative phenotype resembling premature aging. **a**, Increased alopecia and gray fur in *Norad^−/−^* males. Representative 12-month-old male mice shown. **b**, Kyphosis in *Norad^−/−^* mice. Kyphosis severity was scored from 0–3 using an established scheme^49^. Upper right graph shows kyphosis incidence (score ≥ 2), lower right graph shows average kyphosis score at 12 months of age. Representative photographs and x-ray images of 12-month-old mice are shown. **c**, Body weight of 12-month-old *Norad^−/−^* mice with a kyphosis score ≥ 2 compared to randomly-selected *Norad^+/+^* controls (n = 14–16 mice per genotype). **d**, Body fat percentage of mice from panel (c), quantified using NMR. Photographs show representative 12-month-old mice with skin removed to demonstrate loss of fat depots. **e**, Subcutaneous (s.c.) fat thickness in 12-month-old *Norad^−/−^* mice with a kyphosis score ≥ 2 compared to randomly-selected *Norad^+/+^* controls (n = 17 mice per genotype). Representative H&E-stained skin sections shown. **f**, Increased oxidative muscle fibers in 12-month-old *Norad^−/−^* mice. Fiber types in gastrocnemius muscle were grouped into slow twitch oxidative (STO), fast twitch oxidative (FTO), or fast twitch glycolytic (FTG) based on their high, intermediate or low SDH activity (n = 4 mice per genotype). Representative images of SDH histochemistry of gastrocnemius middle zones are shown. **g**, Aging-related changes in the CNS of 12-month-old *Norad^−/−^* mice. Semi-thin sections of spinal cord demonstrate the presence of motor neurons with an accumulation of lipofuscin (arrowhead, upper right panel) or vacuoles (arrowhead, lower right panel) in *Norad^−/−^* mice. **h**, Overall survival of mice of the indicated genotypes over a two-year period. Data are represented as mean ± SD in (**b**)-(**f**), and p values were calculated using log-rank test in (**a**), (**b**), and (**h**) or Student’s t test in (**b**)-(**f**). *p ≤0.05, **p ≤0.01, ****p ≤ 0.0001.

### PUMILIO hyperactivity in *Norad*-deficient mice

Previous work established that *NORAD* is the preferred binding partner of PUM2 in human cells^5^. *NORAD* knockout or knockdown triggers PUMILIO hyperactivity and a consequent loss of genomic stability due to excessive PUMILIO-mediated repression of a set of target mRNAs that are critical for normal mitosis^5,6^. Given the prior demonstration that genomic instability in mice causes aging-associated phenotypes^17,18^, the regulation of PUMILIO activity by this lncRNA, and the resulting effects of PUMILIO hyperactivity, could potentially underlie the phenotype of *Norad*-deficient animals.

To assess whether *Norad* regulates PUMILIO activity in mice, we performed enhanced UV crosslinking immunoprecipitation coupled with high-throughput sequencing (eCLIP)^19^ to assess the transcriptome-wide interactions of PUM2 with target RNAs in *Norad^+/+^* and *Norad^−/−^* mice (**Supplementary Table 1**). Brain was chosen for these experiments because *Norad* shows the highest expression in this tissue (Fig. 1b and Supplementary Fig. 1b), pathologic changes are present in the CNS of *Norad^−/−^* mice (Fig. 2g and Supplementary Fig. 2e, f), and mammalian PUMILIO proteins have been implicated in various neuronal functions^10,11,20–24^.

Like human *NORAD*, the mouse transcript is highly enriched for PREs, harboring 11 perfect matches to the canonical PRE consensus and an additional 3 PREs conforming to a slightly relaxed consensus sequence (UGUANAUN) (Fig. 3a). Accordingly, robust binding of PUM2 to *Norad* was detectable by eCLIP, with the majority of binding occurring in the vicinity of PRE sequences. Strikingly, *Norad* was by far the most highly bound PUM2 target in the transcriptome (Fig. 3b), exhibiting at least 1000 times greater CLIP signal than 95% of all PUM2-bound mRNAs. Thus, as observed in human cell lines, but to a much greater extent *in vivo*, *Norad* is the preferred RNA target of PUM2 in mouse brain.

**Fig. 3.**
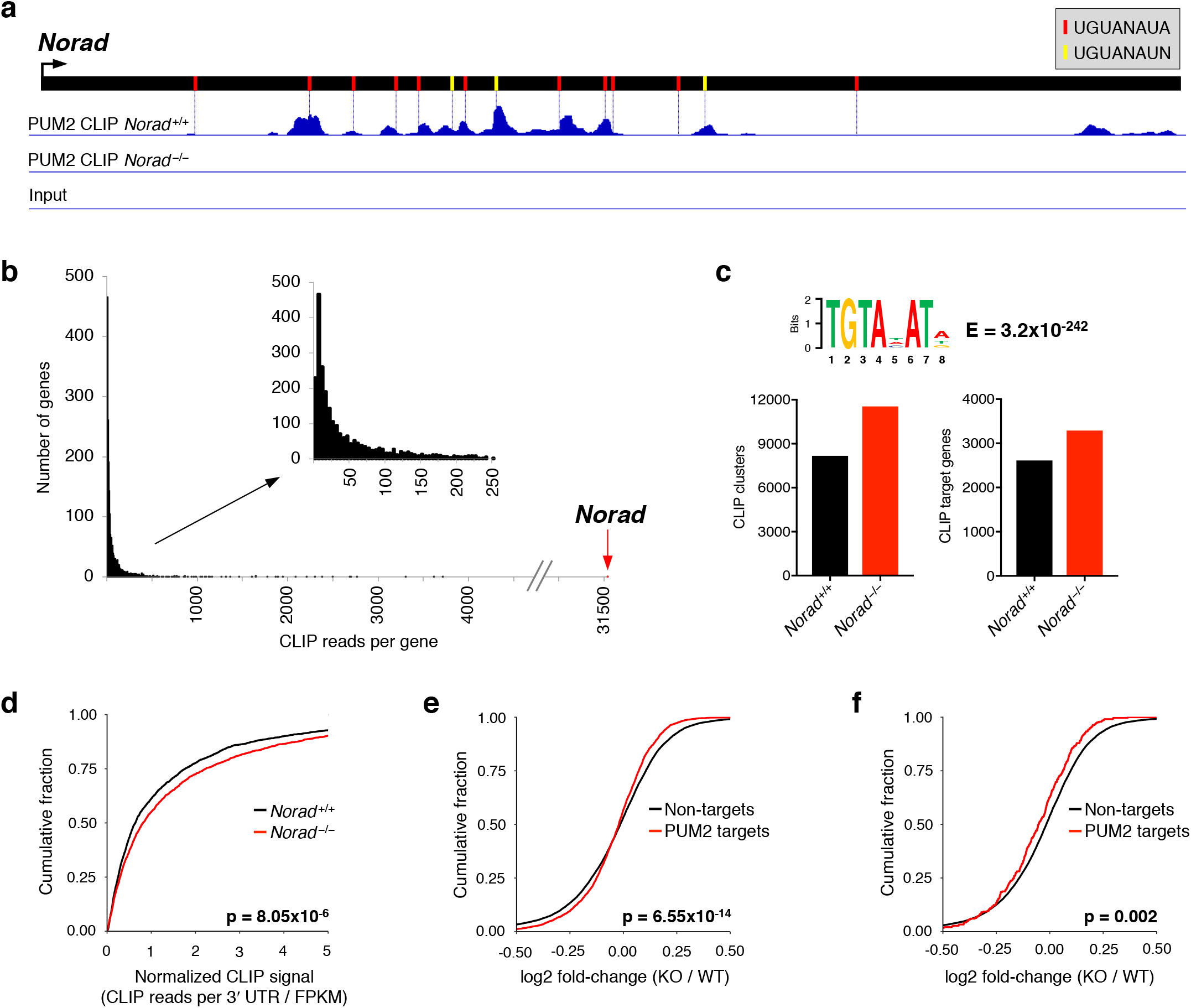
PUMILIO hyperactivity in *Norad*-deficient mice. **a**, Normalized brain PUM2 CLIP reads mapped to *Norad* visualized using the Integrative Genomics Viewer (scale = 0–1260). Positions of perfect or relaxed PREs indicated with red or yellow lines, respectively. **b**, *Norad* is the preferred PUM2 target RNA in mouse brain. The sum of all normalized reads in CLIP clusters in the 3¢ UTR of each PUM2 target RNA was calculated and data were plotted as a histogram showing numbers of genes with a given number of total CLIP reads. **c**, Graphs showing number of 3¢ UTR PUM2 CLIP clusters and number of PUM2 target genes with at least one CLIP cluster in the 3¢ UTR in brains from mice of the indicated genotypes. The web logo above the graphs shows the most significantly enriched motif identified by MEME-ChIP analysis^50^ in CLIP clusters in 3¢ UTRs. **d**, Increased PUM2 target occupancy in *Norad^−/−^* brains. Cumulative distribution function (CDF) plot showing the normalized CLIP signal (total CLIP reads in clusters in the 3¢ UTR normalized to the RNA-seq-determined expression level of the gene) for PUM2 target genes detected in both *Norad^+/+^* and *Norad^−/−^* brains. **e**, CDF plot comparing fold-changes of PUM2 CLIP targets detected in *Norad^−/−^* brains to non-targets. Fold-changes calculated using *Norad^−/−^* vs. *Norad^+/+^* brain RNA-seq data. **f**, As in (e) but only PUM2 CLIP targets with at least a 2-fold increase in normalized CLIP signal in *Norad^−/−^* vs. *Norad^+/+^* brain were plotted. p values calculated by Kolmogorov-Smirnov test for (**d**)-(**f**).

We next examined the transcriptome-wide interactions of PUM2 with target mRNAs. Notably, the relaxed PRE consensus was the most enriched sequence motif detected in PUM2-bound mRNA 3’ UTRs, supporting the reliability of this eCLIP dataset (Fig. 3c). Moreover, PUM2 target occupancy was significantly increased and expression of PUM2 CLIP targets was significantly decreased in *Norad^−/−^* brain, consistent with PUMILIO hyperactivity (Fig. 3d, e). Augmented repression of PUM2 CLIP targets was even more apparent when specifically examining targets whose PUM2 binding was measurably increased in *Norad^−/−^* brains (Fig. 3f). Overall, these data strongly support a conserved function for *Norad* as a negative regulator of PUMILIO activity *in vivo*.

### *Norad* deficiency leads to genomic instability

To assess whether *Norad* loss of function results in genomic instability, fluorescence *in situ* hybridization (FISH) was used to quantify the number of marker chromosomes in primary hematopoietic cells, a representative mitotic tissue. This analysis revealed a significant increase in aneuploid lymphocytes and splenocytes in 3-month and 12-month old *Norad^−/−^* mice (Fig. 4a). To directly determine whether loss of *Norad* results in mitotic abnormalities, murine embryonic fibroblasts (MEFs) were examined using DNA FISH and live cell imaging. In contrast to lymphocytes or splenocytes, we observed a high rate of polyploidization in all MEF lines tested, as previously reported^25^ (Supplementary Fig. 3a, b). We therefore excluded tetraploid and octaploid cells from those scored as aneuploid (higher ploidy was rarely observed). Despite applying these stringent criteria, we detected a significant increase in aneuploidy in *Norad^−/−^* MEFs (Supplementary Fig. 3a). Moreover, time lapse microscopy revealed an increased occurrence of anaphase bridges and lagging chromosomes as *Norad^−/−^* MEFs underwent mitosis (Fig. 4b). Consistent with these findings and our previous observations in *NORAD*-deficient human cell lines^5^, RNA-seq revealed significant repression of genes involved in the cell cycle, mitosis, DNA replication, and DNA repair in *Norad^−/−^* spleens (Fig. 4c, d). These results establish a conserved, essential role for the *Norad*-PUMILIO axis in the maintenance of genomic stability in humans and mice.

**Fig. 4.**
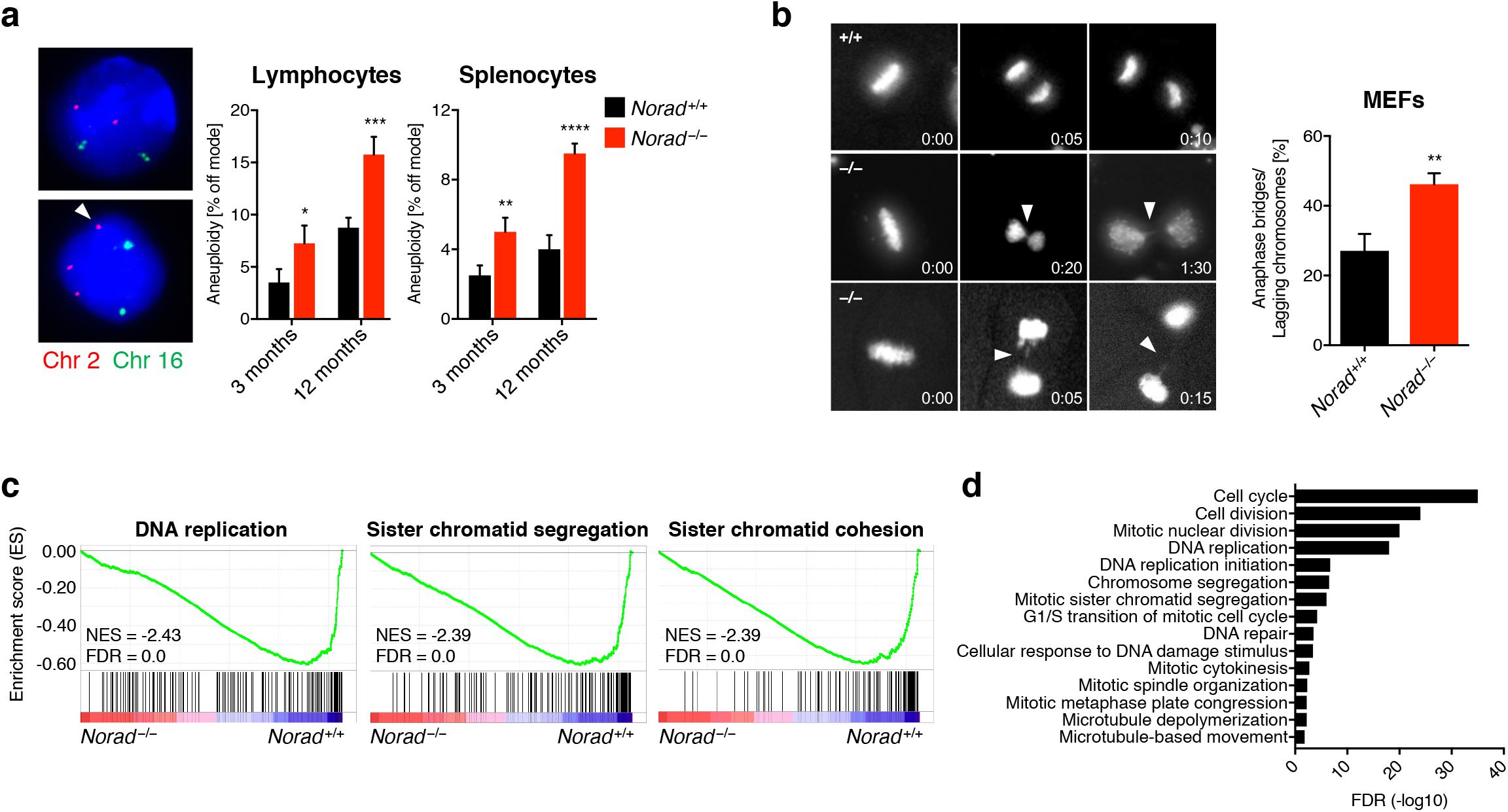
*Norad* deficiency leads to genomic instability. **a**, DNA FISH for two representative chromosomes (chr. 2 and 16) was performed on cultured lymphocytes or freshly-isolated splenocytes from mice of the indicated genotypes and ages. Frequency of cells with a non-modal number of chromosomes was determined by scoring 100 interphase nuclei per sample (n = 4 mice per genotype per time-point). Images show examples of on mode (upper panel) or off mode (lower panel) cells. **b**, Representative time-lapse images of metaphase-to-anaphase transitions of Hoechst-stained primary MEFs show anaphase bridges and lagging chromosomes (arrowheads). Time stamp indicates hours:minutes elapsed. Graph represents data from a total of 128 *Norad^+/+^* and 158 *Norad^−/−^* mitoses analyzed in 3 independent MEF lines per genotype. **c**, GSEA showing repression of indicated gene ontology (GO) gene sets in RNA-seq data from *Norad^−/−^* spleens. **d**, GO analysis was performed using genes that were significantly downregulated in *Norad^−/−^* spleens (EdgeR p ≤ 0.05) using DAVID. Significantly enriched biological processes (BP) are depicted in the graph. Data are represented as mean ± SD in (**a**) and (**b**), and p values were calculated using Student’s t test. *p ≤0.05, **p ≤0.01, ***p ≤0.001, ****p ≤0.0001.

### Loss of *Norad* results in mitochondrial dysfunction

Phenotypic analyses of *Norad*-deficient mice unexpectedly revealed that in addition to genomic instability, widespread mitochondrial dysfunction was evident in knockout tissues. Overt mitochondrial abnormalities were observed in skeletal muscle of 12-month-old *Norad^−/−^* mice, including large accumulations of subsarcolemmal mitochondria (Fig. 5a,b) accompanied by a significant increase in mitochondrial DNA (mtDNA) content (Supplementary Fig. 4a). Ultrastructurally, these mitochondria appeared irregular in shape and enlarged, with loss of cristae (Fig. 5b). Similarly irregular and enlarged mitochondria were observed in spinal neurons of *Norad^−/−^* mice (Fig. 5c). These structural abnormalities were accompanied by evidence of reduced mitochondrial function, such as decreased cytochrome c oxidase (COX; also known as Complex IV of the electron transport chain) activity in spinal neurons (Fig. 5d). Additionally, rare COX-negative fibers were observed in *Norad*-deficient but not wild-type skeletal muscle (Supplementary Fig. 4b). These findings were noteworthy given the extensive evidence linking a decrease in mitochondrial function to aging-associated phenotypes^26^ and the previous demonstration that mice lacking proofreading activity of the mtDNA polymerase, which consequently accumulate mtDNA mutations and deletions, exhibit a premature aging phenotype with many similarities to that seen in *Norad^−/−^* mice^27^.

**Fig. 5.**
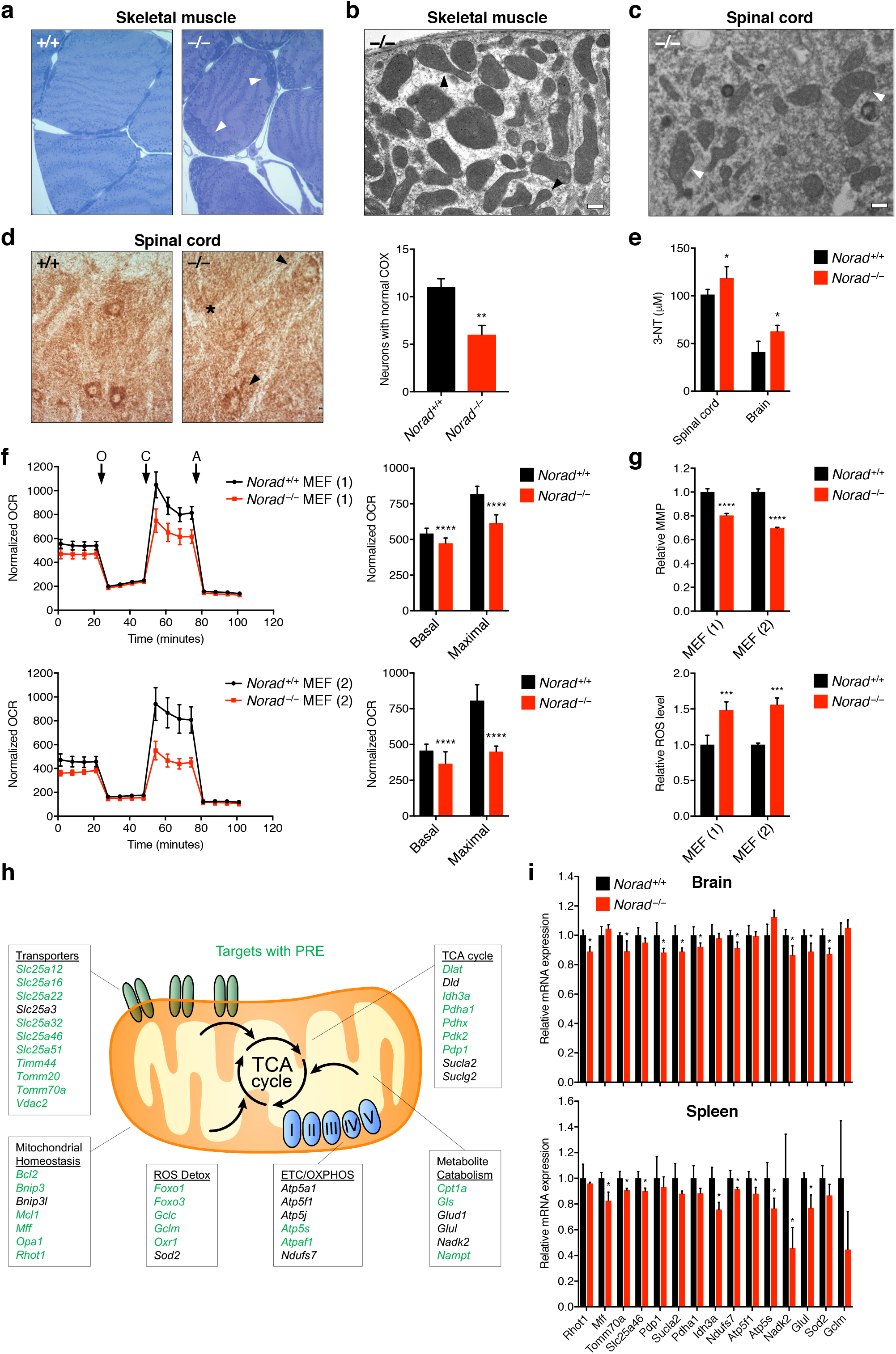
Loss of *Norad* results in mitochondrial dysfunction. **a**, Subsarcolemmal accumulation of mitochondria (arrowheads) in skeletal muscle of 12-month-old *Norad^−/−^* mice. Semi-thin sections of soleus muscle were stained with toluidine blue. **b-c**, Electron micrographs of soleus muscle (**b**) and spinal motor neurons (**c**) showing abnormally enlarged mitochondria without recognizable cristae (arrowheads) in 12-month-old *Norad^−/−^* mice. Scale bars 500 nm. **d**, Reduced COX activity in the CNS of 12-month-old *Norad^−/−^* mice. Spinal cord motor neurons were analyzed using COX histochemistry, and neurons with normal COX activity were counted (n = 7–8 sections total from 4 mice per genotype). Representative images are shown with arrowhead and asterisk pointing to neurons with decreased or absent COX activity, respectively. **e**, Increased protein oxidation in the CNS of 12-month-old *Norad^−/−^* mice, as determined by 3-nitrotyrosine (3-NT) ELISA (n = 3 mice per genotype for brain, n = 4 mice per genotype for spinal cord). **f**, Impaired respiration in immortalized *Norad^−/−^* MEFs. Normalized oxygen consumption rates (OCR) (OCR/total protein) were determined in two littermate pairs of *Norad^+/+^* and *Norad^−/−^* MEFs using Seahorse analysis (n = 22–24 biological replicates per MEF pair, O = oligomycin, C = CCCP, A = antimycin A). Basal and maximal respiration was determined from measurement 4 and 12, respectively. **g**, Reduced mitochondrial membrane potential (MMP) and elevated ROS levels in immortalized *Norad^−/−^* MEFs. MMP and ROS levels were assessed in two littermate pairs of *Norad^+/+^* and *Norad^−/−^* MEFs using flow cytometry (n = 3 biological replicates per MEF pair). **h**, Selected PUM2 CLIP targets with important mitochondrial functions. Green text indicates the presence of a relaxed PRE (UGUANAUN) within 100 nucleotides of a 3¢ UTR PUM2 CLIP cluster. **i**, Downregulation of mitochondrial PUM2 target genes in *Norad^−/−^* brain and spleen. Expression levels were determined by qRT-PCR (n = 4 mice per genotype). Data are represented as mean ± SD in (**d**)-(**g**) and (**i**), and p values were calculated using Student’s t test. *p ≤0.05, **p ≤0.01, ***p ≤0.001, ****p ≤0.0001.

A major consequence of mitochondrial dysfunction that is believed to play a role in cellular damage and aging is the accumulation of reactive oxygen species (ROS)^28,29^. Indeed, brain and spinal cord of *Norad^−/−^* mice show evidence of oxidative damage, including elevated levels of 3-nitrotyrosine (3-NT), 4-hydroxynonenal (4-HNE), and 8-hydroxy-2’-deoxyguanosine/8-hydroxyguanosine (8-OHdG/8-OHG), markers of protein, lipid, and nucleic acid oxidation, respectively (Fig. 5e and Supplementary Fig. 4c).

To directly assess mitochondrial function in *Norad*-deficient cells, respiration rates were analyzed in pairs of littermate-matched *Norad^+/+^* and *Norad^−/−^* MEFs. Basal and maximal respiration was significantly reduced in *Norad^−/−^* cells (Fig. 5f), accompanied by a decrease in mitochondrial membrane potential (MMP) and an increase in ROS production (Fig. 5g). Respiration was also examined in human *NORAD^−/−^* HCT116 cells^5^. Unlike MEFs, HCT116 cells lacking *NORAD* exhibited a significant increase in mitochondrial content (Supplementary Fig. 5a-c). Nevertheless, when normalized to mtDNA copy number, a similar reduction in respiration and increase in ROS was detectable in these cells (Supplementary Fig. 5d, e). These results document a previously unrecognized requirement for *Norad* in the maintenance of mitochondrial homeostasis in mammalian cells and tissues.

To investigate the mechanism through which *Norad* loss-of-function leads to mitochondrial dysfunction, we examined RNA-seq data from *Norad^+/+^* and *Norad^−/−^* brain and spleen using Gene Set Enrichment Analysis (GSEA)^30^. Genes associated with mitochondria-related gene ontology (GO) terms, such as mitochondrial protein complex, electron transport chain, and oxidative phosphorylation, were significantly repressed in *Norad*-deficient tissues (Supplementary Fig. 6a). Remarkably, identical gene sets were repressed in human *NORAD^−/−^* HCT116 cells. We further identified a set of PUM2 brain CLIP targets within these gene sets that are known to perform important functions in mitochondrial biogenesis and homeostasis, mitochondrial transport, oxidative phosphorylation, metabolism, and ROS detoxification (Fig. 5h). Downregulation of a representative set of these genes was validated by qRT-PCR in *Norad^−/−^* brain, spleen, and multiple independent MEF lines (Fig. 5i and Supplementary Fig. 6b). These data are consistent with a model in which PUMILIO hyperactivity in *Norad-*deficient cells and tissues leads to coordinated downregulation of a broad set of target genes that are critical for normal mitochondrial function.

### Enforced PUM2 expression phenocopies *Norad* loss of function

While widespread genomic instability and mitochondrial dysfunction would be predicted to result in the premature aging-like phenotype displayed by *Norad^−/−^* mice, it remained to be demonstrated whether PUMILIO hyperactivity alone could account for the full spectrum of observed phenotypes. To address this question, transgenic mice with doxycycline (dox)-inducible expression of FLAG-tagged PUM2 were generated and crossed to mice harboring a ubiquitously-expressed reverse tetracycline-controlled transactivator 3 transgene (*CAG-rtTA3*)^31^ (Fig. 6a). Administration of dox induced broad transgene expression in *Pum2; rtTA3* double transgenic mice, as documented by FLAG immunohistochemistry (IHC) (Supplementary Fig. 7a). Surprisingly, total levels of PUM2 protein were not increased at the bulk tissue level (Supplementary Fig. 7b), although this finding is consistent with the known negative feedback of PUM1 and PUM2 on their own transcripts^32–34^. Thus, these transgenic mice represent a model of deregulated, but not overtly overexpressed, PUM2. Because *CAG-rtTA3* does not efficiently drive transgene expression in the CNS^31^, we focused our phenotypic studies of *Pum2; rtTA3* double transgenic mice on peripheral tissues.

**Fig. 6.**
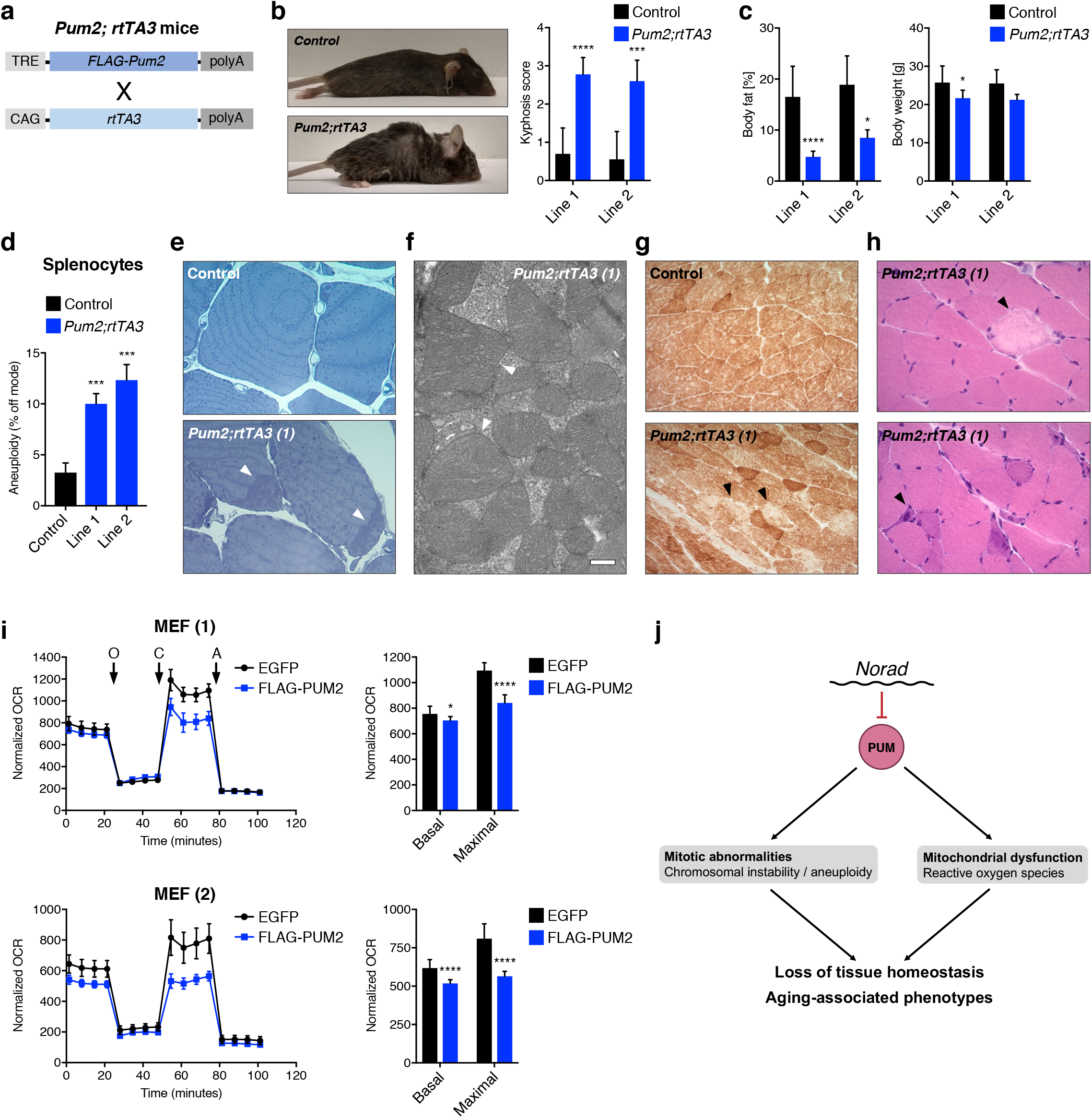
Enforced PUM2 expression phenocopies *Norad* loss of function. **a**, Schematic depicting transgenes in doxycycline (dox)-inducible *Pum2* mice (TRE, TET responsive element; CAG, CAG promoter; rtTA3, reverse tetracycline-controlled transactivator 3; polyA, polyadenylation site). **b**, Kyphosis in *Pum2; rtTA3* mice. Kyphosis scores were determined in two independent transgenic lines after 1.5–2 months of dox treatment. Dox-treated littermate matched wild-type and single transgenic *rtTA3* and *Pum2* mice were used as controls (n = 5–10 mice per genotype per transgenic line). **c**, Reduced body fat and weight in *Pum2; rtTA3* mice after 1.5–2 months of dox treatment. Whole-body fat was quantified using NMR. Controls as in (b). (n = 7–12 (line 1) or 3–5 (line 2) mice per genotype). **d**, Increased aneuploidy in *Pum2; rtTA3* splenocytes. DNA FISH was performed and quantified using freshly-isolated splenocytes after 2 months of dox treatment as in Fig. 4a. Controls represent dox-treated single transgenic *rtTA3* and *Pum2* mice from line 1. 100 interphase nuclei per mouse were scored (n = 3–4 mice per genotype). **e**, Subsarcolemmal accumulation of mitochondria (arrowheads) in skeletal muscle of *Pum2; rtTA3* (line 1) mice after 1.5 months of dox treatment. Representative dox-treated *Pum2* single transgenic control shown. Semi-thin sections of soleus muscle were stained with toluidine blue. **f**, Representative electron micrograph showing abnormally enlarged mitochondria with distorted cristae (arrowheads) in skeletal muscle (soleus) of dox-treated *Pum2; rtTA3* (line 1) mice. Scale bar 500 nm. **g**, COX histochemistry showing reduced COX activity in gastrocnemius muscle from a dox-treated *Pum2; rtTA3* (line 1) mouse compared to littermate matched dox-treated *Pum2* single transgenic control. Arrowheads indicate muscle fibers with extreme reduction in COX activity. **h**, Histologic analysis of H&E-stained sections of gastrocnemius muscle showing necrotic fibers (upper panel, arrowhead) and basophilic regenerating fibers (lower panel, arrowhead) in dox-treated *Pum2; rtTA3* (line 1) mice. **i**, Impaired respiration in immortalized MEFs transiently overexpressing FLAG-PUM2. Normalized oxygen consumption rates (OCR) (OCR/total protein) were analyzed in two different *Norad^+/+^* MEF lines with either FLAG-PUM2 or EGFP overexpression using Seahorse analysis (n = 10–12 biological replicates per MEF line, O = oligomycin, C = CCCP, A = antimycin A). Basal and maximal respiration was determined from measurement 4 and 12, respectively. **j**, Proposed mechanism for the aging-associated phenotypes associated with disruption of the *Norad*-PUMILIO axis. Data are represented as mean ± SD in (**b**)-(**d**) and (**i**), and p values were calculated using Student’s t test. *p ≤0.05, ***p ≤0.001, ****p ≤0.0001.

Administration of dox to young (8–14 week old) *Pum2; rtTA3* double transgenic mice derived from two independent founders, but not to *Pum2* or *rtTA3* single transgenic controls, resulted in a striking phenotype within two months that closely resembled the appearance of *Norad^−/−^* mice at one year of age. Dox-treated *Pum2; rtTA3* mice developed rapidly progressing kyphosis, alopecia, graying of fur, and loss of body fat (Fig. 6b, c and Supplementary Fig. 7c). These phenotypes were accompanied by increased aneuploidy in splenocytes (Fig. 6d) and the accumulation of subsarcolemmal, irregularly shaped mitochondria lacking normal cristae in skeletal muscle (Fig. 6e, f). Further demonstrating mitochondrial abnormalities, a global reduction in COX activity was observed in *Pum2; rtTA3* skeletal muscle (Fig. 6g and Supplementary Fig. 7d) together with scattered necrotic and regenerating fibers (Fig. 6h and Supplementary Fig. 7e).

Lastly, we directly assessed whether enforced PUM2 expression impairs mitochondrial function in MEFs and human cell lines. Transient expression of FLAG-PUM2 in MEFs or stable expression of either PUM1 or PUM2 in HCT116 significantly impaired respiration (Fig. 6i and Supplementary Fig. 7f, 8a-c). Overall, these data provide compelling evidence that PUMILIO hyperactivity in *Norad*-deficient animals results in genomic instability, mitochondrial dysfunction, and ultimately a multi-system degenerative phenotype resembling premature aging.

## DISCUSSION

Although important roles for a growing number of lncRNA-encoding loci have been uncovered in development and disease states^3,4,35^, definitive examples of noncoding RNA-mediated functions that are essential for mammalian physiology and maintenance of homeostasis across tissues are limited. Our studies of the murine *Norad* ortholog reported here unequivocally establish the importance of this lncRNA, and the tight regulation of its target PUMILIO proteins, in mammalian biology and implicate the *Norad*-PUMILIO axis as a major regulator of aging-associated phenotypes (Fig. 6j). These findings provide important new insights and open new lines of investigation into the roles of noncoding RNAs and RNA binding proteins in normal physiology, aging, and disease.

While PUMILIO proteins belong to a deeply conserved family of post-transcriptional regulators, obvious *Norad* orthologs are apparent only in mammals. Other RNAs that regulate the activity of PUMILIO-related proteins in a similar manner in non-mammalian species have not been reported. Why then did this additional layer of PUMILIO regulation evolve? A possible answer to this question may relate to recent findings that, together with those reported here, demonstrate an exquisite sensitivity to PUMILIO dosage in mammals. Zoghbi and colleagues recently showed that slightly reduced *PUM1* dosage causes neurodegeneration in human and mouse brain^10,11^. Human subjects carrying heterozygous *PUM1* deletions or missense mutations develop a neurodevelopmental disorder referred to as *PUM1*-associated developmental disability, ataxia, and seizure (PADDAS), associated with a ~50% reduction in PUM1 protein, or a later onset variant known as *PUM1*-related cerebellar ataxia (PRCA), associated with only a ~25% lowering of PUM1 levels^10^. Taken together with our findings from this study, in which we examined for the first time the effects of mammalian PUMILIO hyperactivity *in vivo*, we can conclude that PUMILIO activity must be maintained within a very narrow range in order to prevent widespread deleterious consequences. In light of these findings, we propose that one major function of *Norad* is to buffer PUMILIO activity such that it stays within this critical range. This model posits the existence of *Norad*-bound and free PUMILIO pools which are exchangeable and in equilibrium, ensuring a consistent amount of available PUMILIO for target mRNA engagement and preventing fluctuations in PUMILIO expression from manifesting in altered target repression. Indeed, an RNA such as *Norad* represents an ideal molecule to serve as a buffer of this type, as it is able to efficiently regulate the activity of a pre-existing pool of PUMILIO at the level of target engagement.

In addition to providing a buffering function, it is likely that *Norad* is also utilized for dynamic regulation of PUMILIO activity under selected conditions. In particular, this lncRNA is known to be induced by a variety of cellular stressors, including DNA damage^5^ and hypoxia^36^, which would be expected to result in de-repression of PUMILIO targets following these stimuli. Although the functional consequences of modulating PUMILIO-mediated gene regulation under these conditions is not yet understood, continued investigation of the signaling inputs that control this system and the resulting effects on PUMILIO-regulated gene networks will be important to further elucidate the roles of this newly-discovered pathway in mammalian biology.

A surprising observation reported in this study was the rapid onset of dramatic premature aging-like phenotypes in *Pum2* transgenic mice despite a lack of overt overexpression of PUM2 protein at the bulk tissue level. Given that production of transgenic FLAG-PUM2 is robustly detectable by IHC, the repression of endogenous PUMILIO through a previously described negative feedback mechanism^32–34^ provides a likely explanation for the lack of increase in total PUM2 levels. This finding nonetheless raises the question of how transgene induction is able to drive such a striking phenotype if the protein product does not accumulate to supraphysiologic levels. An appealing hypothesis to explain this observation postulates that there are key vulnerable cell populations that become dysfunctional or damaged upon *Pum2* induction. These cells may be rare or may naturally express lower levels of PUMILIO, such that a large change in PUM2 expression within them may be masked by the majority of cells in the tissue.

Alternatively, these cells may exhibit dynamic regulation of PUMILIO levels, for example as they transit the cell cycle, which is perturbed by heterologous *Pum2* expression. Cells that are sensitive to enforced *Pum2* expression likely include stem cell populations whose dysfunction could lead to loss of tissue homeostasis and degenerative phenotypes. Accordingly, PUMILIO and related proteins have been implicated as critical stem cell regulators in model organisms and mammals^37–42^. Identification of cell populations that drive aging-associated phenotypes under conditions of *Norad*-deficiency or PUMILIO hyperactivity represents an important priority for future work as this approach may reveal new cell types whose dysfunction contributes to the natural aging-associated decline of tissue homeostasis and renewal.

Analyses of *Norad*-deficiency and enforced *Pum2* expression unexpectedly revealed that PUMILIO hyperactivity triggers the coordinated repression of a large set of PUM2 target transcripts with key roles in mitochondrial function and homeostasis, associated with widespread structural and functional mitochondrial defects. A large body of evidence has linked a decline in mitochondrial function to aging-associated phenotypes^26^, including the direct demonstration that “mitochondrial mutator mice”, which harbor a mutation in the mitochondrial DNA polymerase and consequently accumulate mtDNA mutations, develop a premature aging phenotype with many features in common with *Norad^−/−^* mice^27^. Thus, mitochondrial dysfunction in concert with genomic instability, another abnormality associated with premature aging in mice^17,18^, provides a compelling mechanistic basis for the phenotype of *Norad*-deficient animals. The regulation of mitochondrial biogenesis and function by PUMILIO-related proteins is not restricted to mammals. The budding yeast Puf family member Puf3p preferentially associates with mRNAs that encode mitochondrial proteins and facilitates their local translation in the vicinity of the mitochondrial protein import machinery^43–45^. In *Drosophila* and cultured mammalian cells, PUMILIO proteins repress translation of mRNAs that encode mitochondria-destined proteins until these transcripts are docked at the mitochondrial outer membrane^46^. Together, these observations suggest a deeply conserved role for PUMILIO proteins in the regulation of mitochondrial biology across eukaryotic species.

Perhaps the most intriguing question to arise from these studies is whether dysregulation of the *Norad*-PUMILIO axis plays a role in physiologic aging and/or human disease. Remarkably, a recent RNA-seq study of noncoding RNA expression in the subependymal zone of human brains of increasing age reported a strong age-related decrease in *NORAD* expression^47^. These findings take on added significance in light of our new understanding of the consequences of disruption of the *Norad*-PUMILIO axis and suggest that this lncRNA, and its target PUMILIO proteins, represent new candidates whose altered expression or function may influence the normal age-related decline in tissue function. These genes also represent previously unrecognized candidates that may be mutated or otherwise disrupted in rare progeroid cases that are unlinked to the genes that are presently known to cause these disorders. Thus, further study of the *Norad*-PUMILIO axis, and the pathways that regulate this noncoding RNA and its target proteins, promises to reveal important and unexpected new insights into mammalian physiology and disease.

## ACKNOWLEDGEMENTS

We thank Feng Zhang for plasmids, Jeanetta Marshburn-Wynn for assistance with mouse husbandry, Robert Hammer and the UT Southwestern Transgenic Core for assistance with mouse generation, James Richardson, John Shelton, Cheryl Lewis, the UT Southwestern Histopathology Core, and the UT Southwestern Tissue Management Shared Resource for assistance with histopathology, Vanessa Schmid and the McDermott Center Next Generation Sequencing Core for high-throughput sequencing, Orhan Oz and Xiankai Sun from the UT Southwestern Department of Radiology for mouse imaging, the Experimental Neuromuscular Laboratories in the Center for Gene Therapy at Nationwide Children’s Hospital for technical support, Jose Cabrera for assistance with graphics, and Kathryn O’Donnell, Eric Olson, and members of the Mendell laboratory for helpful comments on the manuscript. This work was supported by grants from CPRIT (RP160249 to J.T.M.; RP150596 for the UTSW Bioinformatics Core Facility), NIH (R35CA197311 to J.T.M.; P30CA142543 to J.T.M. and the UT Southwestern Tissue Resource; and P50CA196516 to J.T.M.), and the Welch Foundation (I-1961–20180324 to J.T.M.). F.K. is supported by the Leopoldina Fellowship Program (LPDS 2014–12) from the German National Academy of Sciences Leopoldina. J.T.M. and H.Y. are Investigators of the Howard Hughes Medical Institute.

## AUTHOR CONTRIBUTIONS

F.K., M.E.Y., S.L., F.A.G., M.M.E., and Z.S. performed experiments. B.C., H.Z., and Y.X. performed bioinformatic analyses. Z.S. and M.E.Y. performed neuromuscular phenotyping while F.K. and J.T.M. performed other aspects of mouse phenotyping. S.S. and H.Y. assisted with live cell imaging. P.M. assisted with mitochondrial studies. F.K. and J.T.M. wrote the manuscript.

## AUTHOR INFORMATION

The authors declare no competing interests. Correspondence and requests for materials should be addressed to.

## METHODS

### Generation of mice with *Norad* deletion or enforced PUM2 expression

All animal protocols were approved by the Institutional Animal Care and Use Committee (IACUC) of The University of Texas Southwestern Medical Center (UTSW) and The Ohio State University, Nationwide Children’s Hospital. Mice were maintained in regular housing with a 12-hour light/dark cycle and normal chow and water *ad libitum*. *Norad^−/−^* mice were generated in the UTSW Transgenic Core by injecting Cas9 mRNA (Sigma-Aldrich) together with two *in vitro* transcribed sgRNAs flanking the *Norad* locus into fertilized C57BL/6J oocytes as described^51^. Founder mice harboring deletions of *Norad* were maintained by backcrossing to wild-type C57BL/6J mice. Of note, *Norad^−/−^* lines were produced from three independent founder mice with *Norad* deletions (Supplementary Fig. 1c). All were phenotypically indistinguishable and used for subsequent studies of *Norad* function.

Doxycycline (dox)-inducible *FLAG-Pum2* transgenic mice (in this study referred to as *Pum2* mice) were generated in a C57BL/6J background by the UTSW Transgenic Core using standard procedures for pronuclear injection. For the generation of the transgene vector, a cDNA clone of isoform 3 of the mouse *Pum2* coding sequence (BC041773) was purchased from transOMIC Technologies, verified by Sanger sequencing, PCR amplified with primers adding a FLAG tag to the protein N-terminus, and cloned into the pTRE-Tight vector (Clontech). Of note, isoform 3 encodes for the shorter PUM2 variant, which can be detected as the lower of two PUM2 bands in western blots. Broad endogenous expression of this isoform was detected by PCR in all tested mouse tissues (data not shown), confirming its physiologic relevance. Transgene positive *Pum2* mice were crossed to a ubiquitous cytomegalovirus early enhancer element chicken beta-actin (CAG) promoter-driven reverse tetracycline-controlled transactivator 3 (*rtTA3*) mouse line, which was generated in the Lowe laboratory^31^ and obtained from The Jackson Laboratory (stock number 016532). The resulting *Pum2; rtTA3* double transgenic mice were used for subsequent experiments together with *Pum2*, *rtTA3,* and wild-type littermates as controls. Transgene expression was induced in 8–14 week-old mice for 1.5–2 months by administering 2 g/L doxycycline hydrochloride (dox) (Sigma-Aldrich) supplemented with 10 g/L sucrose (Research Products International) in drinking water.

### Isolation of lymphocytes and splenocytes

Mice were anesthetized with isoflurane (Henry Schein Animal Health) and subjected to retro-orbital bleeding. Approximately 200 µL of blood were collected, immediately heparinized with 500 USP units/mL (Fresenius Kabi), transferred to 1.3 mL of PB Max Karyotyping Medium supplemented with 50 µg/mL lipopolysaccharide (Gibco and Sigma-Aldrich), and incubated at 37°C for 48 hours with shaking. After this incubation, cells were harvested and processed for DNA FISH analysis. For the isolation of splenocytes, mice were euthanized with an overdose of isoflurane, and spleens were resected, minced with a razor blade in 1X Hank’s balanced salt solution (HBSS) without calcium and magnesium (Gibco), passed through a 70 µm cell strainer (Corning), and washed with 1X phosphate buffered saline (PBS) (Sigma-Aldrich). Immediately after isolation, splenocytes were processed for DNA FISH analysis.

### Generation and culture of mouse embryonic fibroblasts (MEF)

*Norad^+/–^* females were bred to *Norad^+/–^* males and euthanized at embryonic day E14.5. Under sterile conditions, the uterine horns were removed and washed once with 70% ethanol and three times with 1X PBS. The embryos were then released and the heads and all visceral organs removed. The remainder of the embryo was finely minced using razor blades and treated with 0.25% trypsin/EDTA (Gibco) at 37°C for a total of 20 minutes. MEF growth medium consisting of DMEM with 4.5 g/L glucose, L-glutamine and sodium pyruvate (Gibco) supplemented with 1X nonessential amino acids (NEAA), 1X Antibiotic-Antimycotic (AA) (all Gibco), and 10% fetal bovine serum (FBS) (Gibco, Sigma-Aldrich) was added to inactivate the trypsin. Tissue chunks were disrupted by vigorous pipetting, centrifuged, resuspended in MEF growth medium, and plated in T25 flasks. The next day, non-adherent tissue debris was used for genotyping, while attached cells were transferred to a fresh tissue culture dish and designated as primary MEF passage 1 (P1). Aliquots of primary MEF P1 were frozen and stored until needed. Primary MEFs were used for a maximum of 4 passages. Immortalized MEF lines were generated by transfecting primary MEFs with pSG5-SV40-Large-T-Antigen using Lipofectamine 3000 (Invitrogen) according to the manufacturer’s instructions. Starting at 48 hours post transfection, cells were serially passaged 1:10 to select for SV40-immortalized MEFs. After 6 passages, all SV40 large T antigen-transfected MEF lines were regarded as immortalized and this passage was designated as immortalized MEF P1. Of note, the genders of the MEF lines used in this study are not known. All MEF lines were tested and confirmed to be mycoplasma free.

### Generation and culture of human cell lines

The male colon cancer cell line HCT116 was obtained from ATCC (CCL-247) and cultured in McCoy’s 5a medium (Gibco) supplemented with 1X AA (Gibco) and 10% FBS (Gibco, Sigma-Aldrich). The cell line was authenticated by ATCC using short tandem repeat (STR) analysis in November 2017. The generation of HCT116 *NORAD^−/−^* clones via transcription activator-like effector nuclease (TALEN)-mediated insertion of a Lox-Stop-Lox cassette as well as HCT116 PUM1 and PUM2 overexpression clones using lentiviral transduction was described previously^5^. Cell lines were tested and confirmed to be mycoplasma free.

### RNA isolation and quantitative reverse transcription PCR (qRT-PCR)

Total RNA was isolated from cells or tissues using the miRNeasy Mini Kit (Qiagen) following the manufacturer’s instructions including an on-column DNAse I digest to remove genomic DNA contamination. Complementary DNA (cDNA) was generated from 1 µg of total RNA using the SuperScript III First-Strand Synthesis SuperMix for qRT-PCR (Invitrogen) according to the manufacturer’s protocol. Relative *Norad* expression in mouse was quantified using the Applied Biosystems TaqMan assay for *2900097C17Rik* (*Norad*) (Mm04242407_s1) and the TaqMan Universal II Master Mix (Applied Biosystems). Human *NORAD* was quantified using a custom TaqMan assay described previously^5^. For all other genes analyzed in this study, expression was quantified using the Power SYBR Green PCR Master Mix (Applied Biosystems) together with primers provided in **Supplementary Table 2**. RNA expression levels were normalized to 18S ribosomal RNA (cell line studies) or *Gapdh* mRNA (*in vivo* studies) using either standard curves of each gene or the comparative DCt method. The number of biological replicates is stated in the figure legends, each biological replicate was run with 3 technical replicates.

### Next-generation RNA sequencing (RNA-seq)

Total RNA was isolated from brains and spleens of 10-week-old male *Norad^+/+^* and *Norad^−/−^* mice (3 mice per genotype) using the miRNeasy Mini Kit (Qiagen) including a DNAse I digestion step to remove genomic DNA. RNA integrity was determined with the Agilent 2100 Bioanalyzer, and only RNA samples with an RNA integrity number (RIN) of greater than 8 were used for subsequent analysis. Sequencing libraries were prepared with the TruSeq Stranded Total RNA LT Sample Prep Kit (Illumina) and sequenced using the 75 base pair (bp) single-read protocol on a NextSeq 500 platform (Illumina). On average, a single-read depth of 49 million was obtained for each sample. Library prep and RNA-seq were performed by the UTSW McDermott Center Next-Generation Sequencing Core.

Quality assessment of the RNA-seq data was done using the NGS QC Toolkit (v2.3.3)^52^. Reads with more than 30% of nucleotides with a Phred quality score of less than 20 were removed from further analysis. Quality-filtered reads were then aligned to the mouse reference genome GRCm38 (mm10) using Tophat2 (v2.0.12) with default settings^53^. Only reads uniquely mapped to the genome were kept for future analysis. Aligned reads were counted per gene ID using featureCount (v1.4.6)^54^. Differential gene expression analysis was carried out using the R package EdgeR (v3.8.6)^55^. For each comparison, genes were required to have at least 1 read in at least 1 sample to be considered as expressed. Differential gene expression analysis was performed using the GLM approach following EdgeR’s official documentation. CPM (counts per million) and FPKM (fragments per kilobase million) were obtained using EdgeR and Stringtie (v1.2.2)^56^, respectively.

Gene set enrichment analysis (GSEA)^30^ was performed using default gene sets of gene ontology (GO) terms. The results obtained from the EdgeR RNA-seq analyses of brain and spleen (normalized reads as CPM) were used as input data. The normalized enrichment scores (NES) as well as the false discovery rates (FDR) are provided in the figures. GO analysis of spleen RNA-seq data was also carried out on genes that were significantly (p £ 0.05) downregulated in *Norad^−/−^* spleens using the Database for Annotation, Visualization and Integrated Discovery (DAVID)^57,58^.

### Enhanced UV crosslinking immunoprecipitation (eCLIP)

PUM2 RNA interactions in the mouse brain were determined by eCLIP, following a previously published protocol^19^. In brief, brains from 3-month-old *Norad^+/+^* and *Norad^−/−^* females (2 mice per genotype representing 2 biological replicates) were resected and cut in halves. Per sample, one half of a brain was minced with razor blades in 1X ice-cold diethyl pyrocarbonate (DEPC)-treated PBS. The resulting tissue suspensions were UV crosslinked on ice in a Spectrolinker XL-1500 (Spectronics) at 254nm three times at 400mJ/cm^2^. The UV crosslinked tissues were centrifuged, snap-frozen in ethanol/dry ice, and stored at −80°C until needed. For the immunoprecipitation of PUM2, Protein G Dynabeads (Invitrogen) were used together with the same polyclonal goat anti-PUM2 antibody (sc-31535, Santa Cruz) used previously for PUM2 CLIP analysis in human cells^5^. For each genotype, duplicate size-matched input and immunoprecipitation samples were prepared (4 samples per genotype). In contrast to the original protocol^19^, in which libraries were designed for paired-end sequencing, we adapted the RNA and DNA linker sequences for single-read sequencing. For each sample, separate sequencing libraries were generated using a unique modified RiL19-new RNA linker as well as a modified rand103Tr3-new DNA linker and AR17-new reverse transcription primer (sequences provided in **Supplementary Table 2**). PCR library amplification was performed with polyacrylamide gel electrophoresis (PAGE)-purified oligonucleotides containing specific indexes (D501-D504, D701-D703). Single-read sequencing was performed on a NextSeq 500 platform using a NextSeq 500/550 High Output v2 Kit, 75 cycles (Illumina) in the UTSW McDermott Center Next-Generation Sequencing Core.

All adapter sequences were removed using Cutadapt with an e-value set to 0.1. All reads less than 18 nt after adapter trimming were discarded, and the unique molecular identifiers (10 nt randomers) for PCR duplication identification were trimmed and recorded using in-house scripts. Because of the high number of *NORAD* pseudogenes in the human and mouse genome, we followed a similar mapping strategy to that used in our previous study^5^. Reads were first mapped to *Norad* (*2900097C17Rik*) before all remaining reads, which did not align to *Norad*, were mapped to GRCm38 (mm10) using Tophat2 (v2.0.12) with default settings^53^. Only uniquely mapped genomic reads were retained. PCR duplicates were then removed based on the unique molecular identifier information using in-house scripts. All remaining reads were regarded as usable reads and subjected to cluster calling.

For each IP sample, the read coverage of each nucleotide was calculated and all regions with coverages of equal or greater than 3 were kept as candidate bins. The read counts of each IP/input pair were obtained for every bin, requiring at least a 50% sequence overlap. The fold-changes of the normalized read counts were then calculated for each bin: normalized fold-change = ((reads_in_bin[IP] + 1)/total_usable_reads[IP])/((reads_in_bin[input] + 1)/total_usable_reads[input]). Bins with fold-changes greater than or equal to 4 were considered as clusters. Finally, we filtered clusters for those that were detected in both biological replicates of either genotype. Clusters that overlapped with at least 30% of their length were merged. To annotate PUMILIO response elements (PREs) in CLIP clusters, all final clusters were extended by 100 nt at both ends and analyzed for the occurrence of perfect (TGTANATA) and relaxed (TGTANATN) PREs. Bedgraph files of each sample were generated with BEDtools^59^ using reads normalized to the total usable genomic read count and visualized with the Integrative Genomics Viewer (IGV). Genes with one or more CLIP clusters in their 3¢ UTRs were regarded as PUM2 target genes. The number of CLIP reads per gene was determined by calculating the weighted sum of all reads within 3¢ UTR CLIP clusters for each PUM2 target gene from both *Norad^+/+^* CLIP replicates. To calculate normalized CLIP signal of PUM2 targets in *Norad^+/+^* and *Norad^−/−^* brains, genes were first filtered for those with an average expression level of at least 1 FPKM in brain RNA-seq data. The average normalized number of CLIP reads from both CLIP replicates within 3¢ UTR clusters of each PUM2 target gene were then summed and divided by the gene’s expression level (average FPKM) in the respective genotype.

### Quantification of mitochondrial DNA

Total DNA was isolated using the DNeasy Blood and Tissue Kit (Qiagen) according to the manufacturer’s instructions. Mitochondrial (mtDNA) and nuclear (nDNA) DNA were quantified by qPCR using either human or mouse specific primers (**Supplementary Table 2**) and the Power SYBR Green PCR Master Mix (Applied Biosystems). The quantity of mtDNA was then normalized to the quantity of nDNA. Both mtDNA and nDNA concentrations were determined using standard curves. The number of replicates is provided in the respective figure legends.

### Subcellular fractionation

Immortalized *Norad^+/+^* MEFs were seeded in triplicate and harvested the next day for subcellular fractionation, which was performed as previously described^5^. Briefly, cell pellets were lysed in RLN1 buffer (50 mM Tris-HCl pH 8.0, 140 mM NaCl, 1.5 mM MgCl2, 0.5% NP-40, RNAse inhibitor), incubated on ice for 5 minutes, and centrifuged. The supernatant contained the cytoplasmic fraction, while the pellet contained the nuclear soluble and chromatin fraction. The pellet was lysed again with RLN2 buffer (50 mM Tris-HCl pH 8.0, 500 mM NaCl, 1.5 mM MgCl2, 0.5% NP-40, RNAse inhibitor), incubated on ice for 5 minutes, and centrifuged. The supernatant represented the nuclear soluble fraction, while the pellet yielded the chromatin fraction. All fractions were then subjected to RNA isolation and subsequent qRT-PCR. All samples were tested for *Norad* as well as for *Neat1* (nuclear control) and *Actb* (cytoplasmic control).

### Western blots

Cell and tissue lysates were prepared in RIPA buffer (50 mM Tris-HCl pH 8.0, 150 mM NaCl, 1% NP-40, 0.5% sodium deoxycholate, 0.1% sodium dodecyl sulfate) supplemented with cOmplete Protease Inhibitor Cocktail (Roche). Western blots were probed with monoclonal rabbit anti-PUM2 antibody (ab92390, Abcam), monoclonal rabbit anti-PUM1 antibody (ab92545, Abcam), or monoclonal rabbit anti-GAPDH antibody (2118, Cell Signaling). Bands were visualized using an IRDye 800CW donkey anti-rabbit IgG secondary antibody (925–32213, Licor) and an Odyssey CLx Imager (Licor).

### FLAG immunohistochemistry

Tissues were harvested from mice that had been treated with dox for 3.5–6.5 weeks. All tissues were fixed in 10% neutral buffered formalin (NBF) for 24–48 hours. Fixed samples were processed, paraffin embedded, and sectioned using standard procedures. To detect the expression of the *FLAG-Pum2* transgene, immunohistochemistry (IHC) was performed by the UTSW Tissue Management Shared Resource using the monoclonal anti-FLAG M2 antibody (F1804, Sigma-Aldrich). Images were acquired on an AxioObserver Z1 microscope (Zeiss).

### Histologic analysis of skeletal muscle and the central nervous system (CNS)

12-month-old *Norad^+/+^* and *Norad^−/−^* mice were used for semi-thin and ultrastructural analysis. For these studies, one group of mice were given xylazine/ketamine anesthesia and euthanized by cardiac perfusion with 4% paraformaldehyde followed by 5% glutaraldehyde (both in 0.1 M phosphate buffer). Tissue samples from brain and spinal cord were removed under a dissecting microscope. A second group of mice were perfused with 4% paraformaldehyde and their muscles were removed and further fixed *in situ* in 5% glutaraldehyde. These tissues were dissected into small blocks and processed for plastic embedding using standard methods^60^. Thick (1 µm) sections were stained with toluidine blue and selected blocks were sectioned and examined with an electron microscope (Hitachi H7650). Brain and spinal cord segments were placed in 10% NBF and processed for paraffin embedding. Brain, spinal cord, and skeletal muscle were collected from additional mice for cryostat sectioning. Muscle tissues from *Pum2; rtTA3* as well as *Pum2* and *rtTA3* single transgenic littermates were collected and processed as for *Norad^−/−^* mice. All transgenic mice were between 4 and 6 months old and had been treated with dox for 1.5–2 months.

To analyze neuronal cell loss in the ventral horn neuron pools, 5 µm thick, paraffin embedded and hematoxylin and eosin (H&E) stained lumbar spinal cord sections from *Norad^+/+^* (n = 5) and *Norad^−/−^* (n = 5) mice were analyzed. Motor neuron pools in the anterior horn cell areas of both hemicords from each section were included. For each mouse, sections from 1–2 levels were photographed at 20X magnification. Only cell bodies clearly showing a nucleolus on the plane of the section were considered. Equal numbers of spinal cord levels were analyzed in each group. Mean neuronal densities of anterior horn areas per lumbar cord level were calculated.

Succinate dehydrogenase (SDH) enzyme histochemistry was used to assess metabolic fiber type changes in the aging muscle. For this purpose, muscle fiber types were grouped into three categories: slow twitch oxidative (STO), fast twitch oxidative (FTO), and fast twitch glycolytic (FTG). Twelve µm thick cross sections from the gastrocnemius muscles of 12-month-old *Norad^+/+^* and *Norad^−/−^* mice (n = 4 in each group, 2 males and 2 females) were stained for SDH activity. Three images, each representing a distinct zone of the gastrocnemius muscle (a deep zone predominantly composed of STO, an intermediate zone showing a checkerboard appearance of STO, FTO, or FTG, and a superficial zone predominantly composed of FTG fibers), were taken along the midline axis at 20X magnification using an Olympus BX41 microscope. Muscle fiber types were determined and counted based on their dark (STO), intermediate (FTO), or light (FTG) SDH staining as previously reported^61^.

Mitochondrial function was assessed in spinal cord neurons of 12-month-old *Norad^+/+^* and *Norad^−/−^* mice using cytochrome c oxidase (COX) histochemistry. Twelve µm thick frozen lumbar spinal cord tissue sections were mounted onto superfrost glass slides (Thermo Fisher) and dried at room temperature for 1 hour before the COX enzymatic activity assay was performed. Two sections, 100 µm apart, were analyzed from each mouse. The anterior horn areas of each section were photographed at 10X magnification and the number of neurons with normal COX activity were determined. Counts from the hemicord anterior horn cell area with the higher number of neurons with normal COX activity was included in the analysis. Only cell bodies with an obviously visible nucleolus on the plane of the section and with a COX activity clearly above the background of gray matter were considered. The number of neurons with normal COX activity was counted per each section analyzed. COX activity was also qualitatively analyzed in fresh frozen gastrocnemius muscle sections from 12-month-old *Norad^+/+^* and *Norad^−/−^* mice and 4–6 month-old *Pum2; rtTA3* transgenic mice or controls that had been treated with dox for 1.5–2 months.

### DNA fluorescence *in situ* hybridization (FISH)

DNA FISH for two representative chromosomes (chromosome 2 and 16) was performed in lymphocytes, splenocytes, and primary MEFs. Cells were first incubated in hypotonic KCl solution: lymphocytes were incubated in 75 mM KCl at 37°C for 15 min, splenocytes in 75 mM KCl at room temperature for 30 minutes, and MEFs in 0.4% KCl at room temperature for 8 minutes. Subsequently, cells were centrifuged and resuspended in methanol/acetic acid (3:1), washed twice with methanol/acetic acid (3:1), dropped onto Rite-On Micro Slides (Gold Seal Products), air-dried, and either used immediately or stored at −20°C until needed. DNA FISH was performed using the Mouse IDetect Chromosome Point Probes for chromosome 2 (red) and 16 (green) (IDMP1002-R, IDMP1016-1-G, Empire Genomics) following the manufacturer’s protocol. Slides were mounted with ProLong Diamond Antifade Mountant with DAPI (Invitrogen) and analyzed on an AxioObserver Z1 microscope (Zeiss) using the 100X oil objective. Lymphocytes and splenocytes whose chromosome count differed from 2n for at least one of the two tested chromosomes were regarded as aneuploid or off mode. MEFs were only considered aneuploid when their chromosome count differed from 2n or a multiple of 2n in order to account for the increased polyploidy in this cell type.

### Time-lapse microscopy

Primary *Norad^+/+^* and *Norad^−/−^* MEFs (3 MEF lines per genotype, P<4) were grown on Lab-Tek Chambered Coverglass slides (Thermo Fisher) that were coated with poly-L-lysine (Sigma-Aldrich). Prior to the analysis, DNA was visualized by adding 50 ng/mL Hoechst dye (Invitrogen) to the growth medium. Mitoses were monitored by taking fluorescence images every 5 minutes for ~48 hours on a Leica inverted microscope equipped with a temperature and CO2-controlled chamber, a 63X oil objective, an Evolve 512 Delta EMCCD camera, and the MetaMorph Microscopy Automation and Image Analysis Software (Molecular Devices, LLC). Videos were generated from the acquired time-lapse images and analyzed for the occurrence of mitotic defects including anaphase bridges and lagging chromosomes.

### Assessment of aging-associated phenotypes

*Norad^+/+^*, *Norad^+/–^*, and *Norad^−/−^* mice were continuously monitored over a period of 12 months for the onset and progression of kyphosis as well as alopecia and graying of fur. The kyphosis scoring system was adopted from a previously reported strategy^49^. In brief, a kyphosis score of 0 indicates no kyphosis detectable, a score of 1 indicates the presence of mild kyphosis but the mouse is still able to entirely stretch its spine, and scores of 2 and 3 indicate that there is prominent kyphosis at rest which persists in a mild (score of 2) or a severe (score of 3) form even when the mouse stretches its spine.

### Analysis of whole-body fat and subcutaneous adipose tissue

Whole-body fat, as a percentage of body weight, was measured by nuclear magnetic resonance (NMR) in 3-month and 12-month-old *Norad^+/+^* and *Norad^−/−^* mice, or in 4–6 month-old *Pum2; rtTA3* double transgenic mice and control littermates after 1.5–2 months of dox treatment, using a Bruker Minispec mq10. Subcutaneous (s.c.) adipose thickness of 12-month-old *Norad^+/+^* and *Norad^−/−^* mice was determined using standard H&E-stained skin histology. For every skin sample, images were acquired at 5X magnification across the entire length of the section on an AxioObserver Z1 microscope (Zeiss) using the AxioVision 4.8 software (Zeiss). In these images, the thickness of the s.c. adipose tissue was measured at 15 different points using the AxioVision 4.8 software. The average of the 15 measurements was then calculated to obtain the adipose thickness of each mouse.

### Quantification of mitochondrial membrane potential (MMP)

Tetramethylrhodamine ethyl ester (TMRE) (ENZ-52309, Enzo Life Sciences) was used for measuring mitochondrial membrane potential (MMP) in immortalized *Norad^+/+^* and *Norad^−/−^* MEFs. 80 x 10^3^ cells were seeded in triplicate in 6-well plates in regular growth medium and incubated for 16–18 hours until 70%-80% confluent. Cells were then trypsinized with 0.25% trypsin/EDTA (Gibco), pelleted at 300 g, and resuspended in 500 µL of fresh growth medium containing 50 nM TMRE. Samples were then incubated at 37°C in the dark for 30 minutes and analyzed by flow cytometry using a BD Accuri C6 Cytometer (BD Biosciences). The average and standard deviation of the mean fluorescence intensities of the 3 replicates was calculated for each sample, and each *Norad^−/−^* MEF line was compared to its *Norad^+/+^* littermate control line.

### Analysis of reactive oxygen species (ROS) and oxidative damage

The Enzo Total ROS Detection Kit (ENZ-51011, Enzo Life Sciences) was used for detection of ROS levels in immortalized MEFs and human HCT116 cells. 80 × 10^3^ MEFs were plated in triplicate in 6-well plates in regular growth medium and incubated for 16–18 hours. 60 × 10^3^ HCT116 cells were seeded in triplicate in 24-well plates in regular growth medium and also incubated for 16–18 hours. All cells were 70%-80% confluent at the time of the assay. ROS levels were measured according to the manufacturer’s protocol. Briefly, cells were trypsinized with 0.25% trypsin/EDTA (Gibco), pelleted at 300 g, washed once with Enzo 1X Wash Buffer, and resuspended in 300 µL of freshly prepared Enzo ROS Detection Solution (1 µL ROS dye in 5 mL Wash Buffer). Samples were incubated in the dark for 30 minutes and analyzed by flow cytometry using a BD Accuri C6 Cytometer (BD Biosciences). The average and standard deviation of the mean fluorescence intensities of the 3 replicates was calculated for each sample. Each *Norad^−/−^* MEF line was compared to its *Norad^+/+^* littermate control line.

Oxidative damage was examined in the CNS of 12-month-old *Norad^+/+^* and *Norad^−/−^* mice using IHC and immunofluorescence. Lipid peroxidation was examined in formalin-fixed paraffin embedded spinal cord sections by IHC using the polyclonal rabbit anti-4-Hydroxynonenal (4-HNE) antibody (ab46545, Abcam). Nucleic acid oxidation was assessed in fresh frozen brain sections by immunofluorescence using the mouse monoclonal anti-DNA/RNA Damage antibody (ab62623, Abcam), which detects 8-OHdG/8-OHG. In addition, protein oxidation was assessed in spinal cord and brain using an enzyme-linked immunosorbent assay (ELISA) for 3-nitrotyrosine (3-NT) (MBS262795, Mybiosource) according to the manufacturer’s instructions.

### Seahorse analysis

Oxygen consumption rates (OCRs) were analyzed in immortalized MEFs and human HCT116 cells using a Seahorse Bioscience XF96 Extracellular Flux Analyzer (Seahorse Bioscience/Agilent). 10 × 10^3^ cells per well were plated in Seahorse XF96 cell culture microplates (Agilent) in regular growth medium and incubated for 14–16 hours. For HCT116 cells, Seahorse XF96 Cell Culture Microplates were coated with poly-L-lysine (Sigma-Aldrich) to improve cell attachment. Prior to measurement, cells were equilibrated for 1 hour in Seahorse assay medium (D5030, Sigma-Aldrich) supplemented with 10 mM glucose (Sigma-Aldrich), 1 mM sodium pyruvate (Gibco), and 2 mM L-glutamine (Sigma-Aldrich). OCRs were monitored before and after adding the following mitochondrial inhibitors: 2 µM oligomycin (complex V inhibitor), 1 µM carbonyl cyanide 3-chlorophenylhydrazone (CCCP, uncoupler of oxidative phosphorylation), and 1 µM antimycin A (complex III inhibitor) (all Sigma-Aldrich). OCRs were normalized to the amount of protein in each sample using a bicinchoninic acid (BCA) assay (Thermo Fisher) according to the manufacturer’s instructions. For HCT116 cells, OCRs were further normalized to the mtDNA content to account for differences in mitochondrial content.

### FLAG-PUM2 overexpression in MEFs

To overexpress PUM2 in MEFs, the same *FLAG-Pum2* cDNA that was used for generating the transgenic mouse was cloned into a pBROAD3 vector (Invivogen), in which the Rosa26 promoter was replaced by a strong CAG promoter (pCAG-*FLAG-Pum2*). A pcDNA3-EGFP vector was used for control transfections. FLAG-PUM2 and EGFP were transfected into immortalized *Norad^+/+^* MEFs using 2.5 µg of plasmid DNA and Lipofectamine 3000 (Invitrogen) according to the manufacturer’s protocol. In brief, 100 × 10^3^ cells were seeded into 6-well plates and incubated overnight. The next day, cells were transfected with either pCAG-*FLAG-Pum2* or pcDNA3-EGFP and incubated again overnight. The following day, cells were collected and re-plated for a second transfection, performed identically. After overnight incubation, cells were seeded for Seahorse analysis, as described above. PUM2 overexpression was assessed after the second transfection using western blot.

### Statistical analysis

A comprehensive description of the RNA-seq and eCLIP analysis including the use of software is provided in the respective sections. The significance of the cumulative distribution functions was calculated using the Kolmogorov-Smirnov test and plotted in R. For all other analyses, statistical significance was analyzed using Prism 7 (GraphPad Software). Student’s t tests or log-rank tests (for survival and phenotype incidence) were used to determine statistical significance, which is indicated as *p ≤ 0.05, **p ≤ 0.01, ***p ≤ 0.001, ****p ≤ 0.0001. Data are presented as mean ± SD in all figures except Supplementary Fig. 5d, 8c (left graph) where the data are presented as mean ± SEM. The numbers of replicates are stated in the figure legends.

---

**Supplementary Fig. 1.**
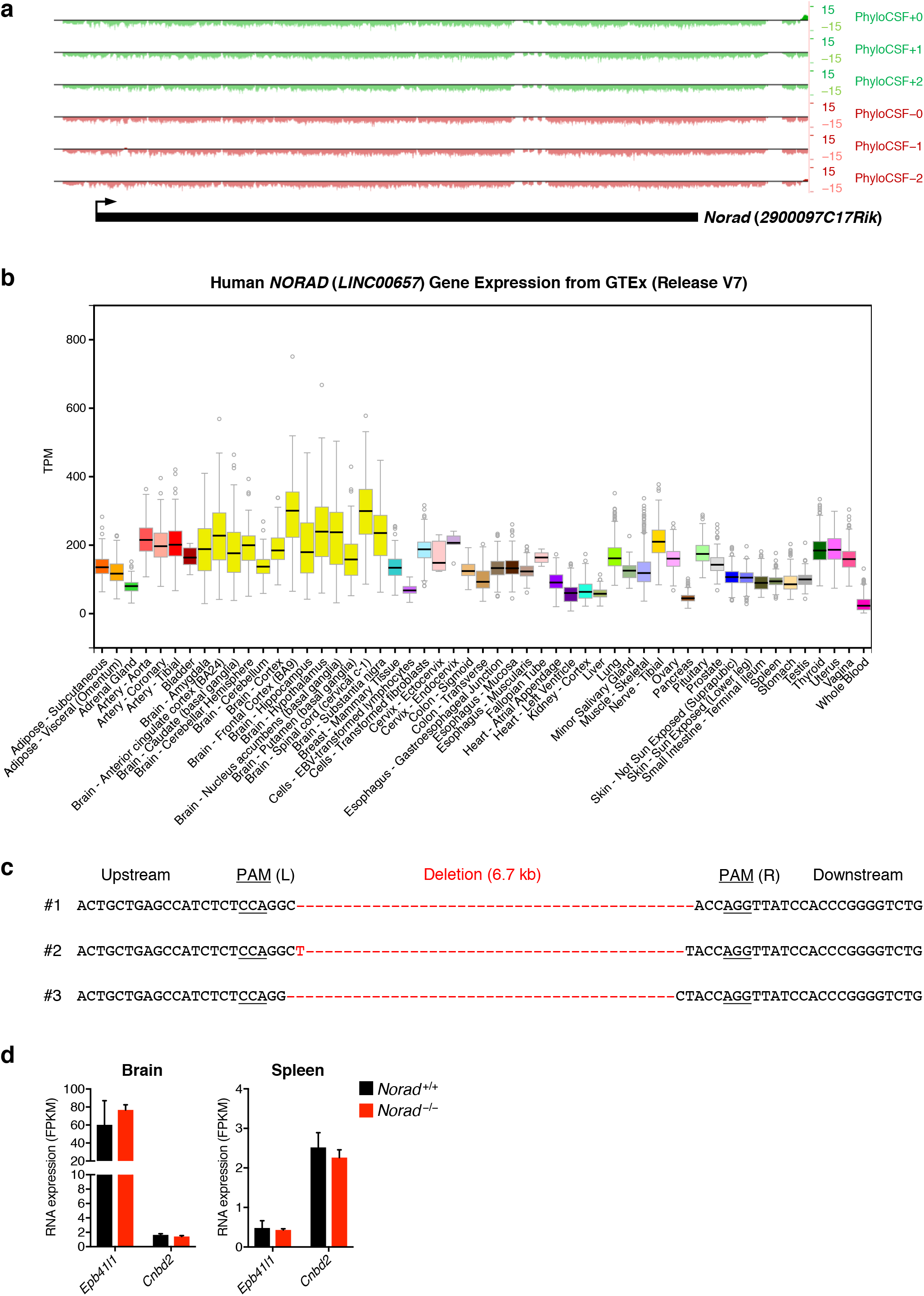
Deletion of the mouse *Norad* ortholog. **a**, Low protein-coding potential of *Norad* (*2900097C17Rik*). PhyloCSF tracks from the UCSC Genome Browser (mm10) shown for all three reading frames on both strands. **b**, *NORAD* (*LINC00657*) expression in human tissues. RNA-seq data from the Genotype-Tissue Expression (GTEx) Project (Release V7) plotted as transcripts per million (TPM). **c**, Sequences of genomic deletions induced by genome editing in *Norad^-/-^* mice. **d**, Expression levels of the indicated genes neighboring *Norad* in brain and spleen as determined by RNA-seq (n = 3 mice per genotype). Data are represented as mean ± SD in (**d**).

**Supplementary Fig. 2.**
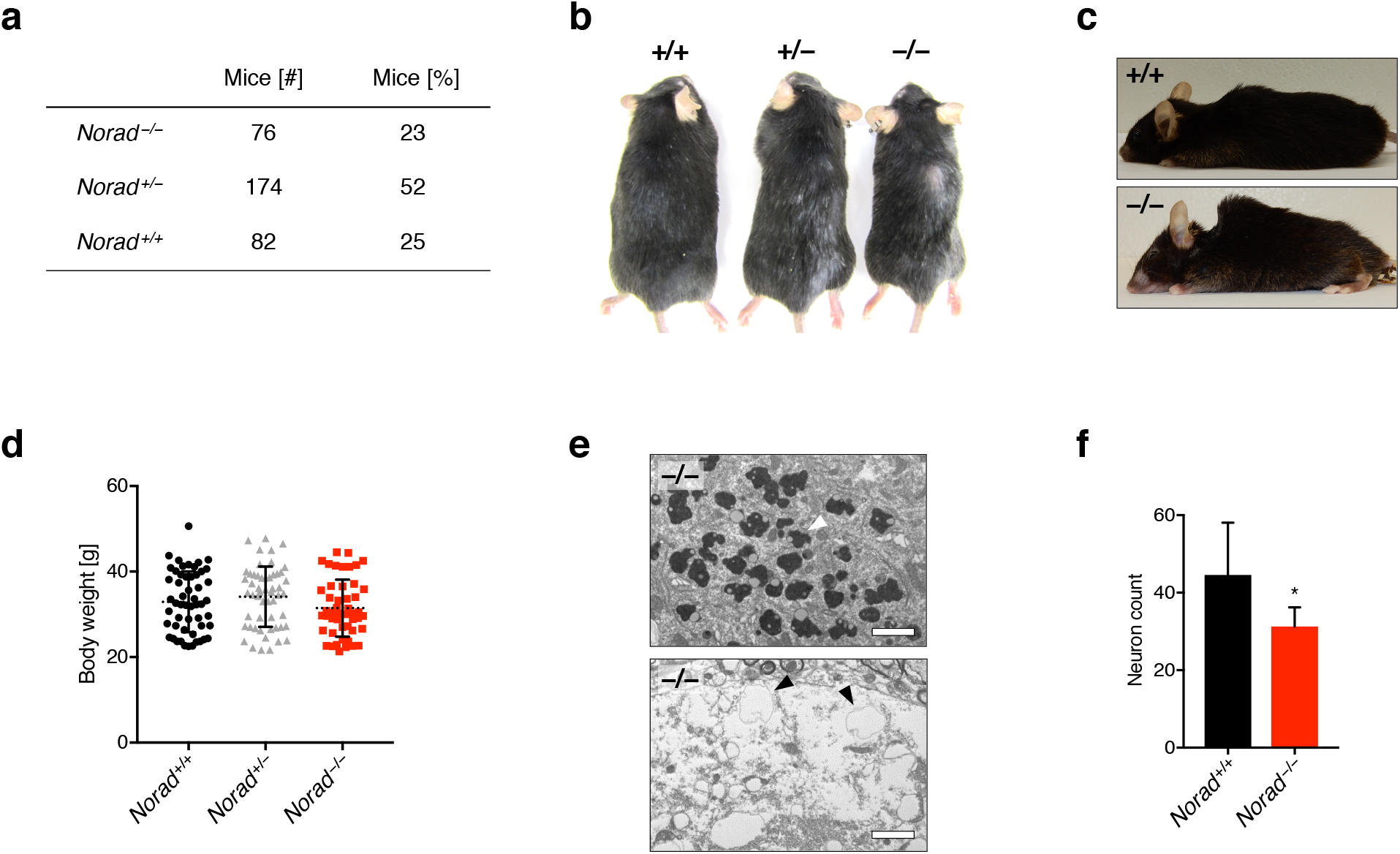
*Norad* loss-of-function results in a degenerative phenotype resembling premature aging. **a**, Distribution of genotypes obtained from *Norad^+/-^* intercrosses. **b**, Representative 12-month-old male *Norad^+/+^, Norad^+/-^* and *Norac^-/-^* littermates showing dose-dependent alopecia and graying of fur. **c**, Pronounced kyphosis in a 34-month-old *Norad^-/-^* mouse compared to an age-matched wild-type control. **d**, Body weight of 12-month-old mice of the indicated genotypes independent of the kyphosis score (n = 48–54 mice per genotype). **e**, Electron micrographs of spinal cord motor neurons in 12-month-old *Norad^-/-^* mice showing an accumulation of lipofuscin (arrowhead, upper panel) and vacuoles (arrowhead, lower panel). Scale bars 2 μm. **f**, Reduction of neuronal density in spinal cord of 12-month-old *Norad^-/-^* mice. For each genotype, neurons were quantified in a total of 7 sections derived from 5 independent mice. Data are represented as mean ± SD in (d) and (f) and p value was calculated using Student’s t test. *p ≤ 0.05.

**Supplementary Fig. 3.**
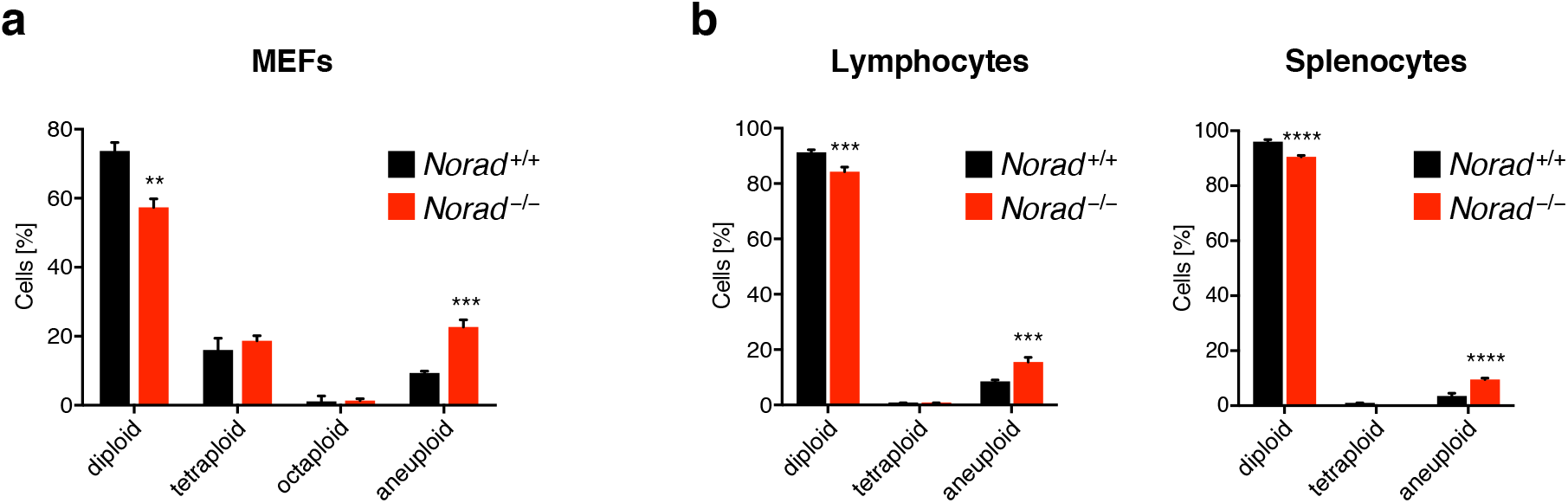
*Norad* deficiency leads to genomic instability. **a**, Aneuploidy and polyploidy in primary MEFs. DNA FISH for two representative chromosomes (chr. 2 and 16) was performed to determine the frequency of diploid, polyploid, and aneuploid cells. Only cells with chromosome numbers differing from 2n or a multiple of 2n were regarded as aneuploid. 120 interphase nuclei per cell line were scored (n = 3 independent MEF lines per genotype). **b**, Rare polyploidy in lymphocytes and splenocytes from 12-month-old *Norad^-/-^* mice. Additional data from the experiment in Fig. 4a is shown (12-month time-point). In only very rare cases (≤ 0.5%) were cells tetraploid for the two tested chromosomes. Data are represented as mean ± SD and p values were calculated using Student’s t test. **p ≤ 0.01, ***p ≤ 0.001, ****p ≤ 0.0001.

**Supplementary Fig. 4.**
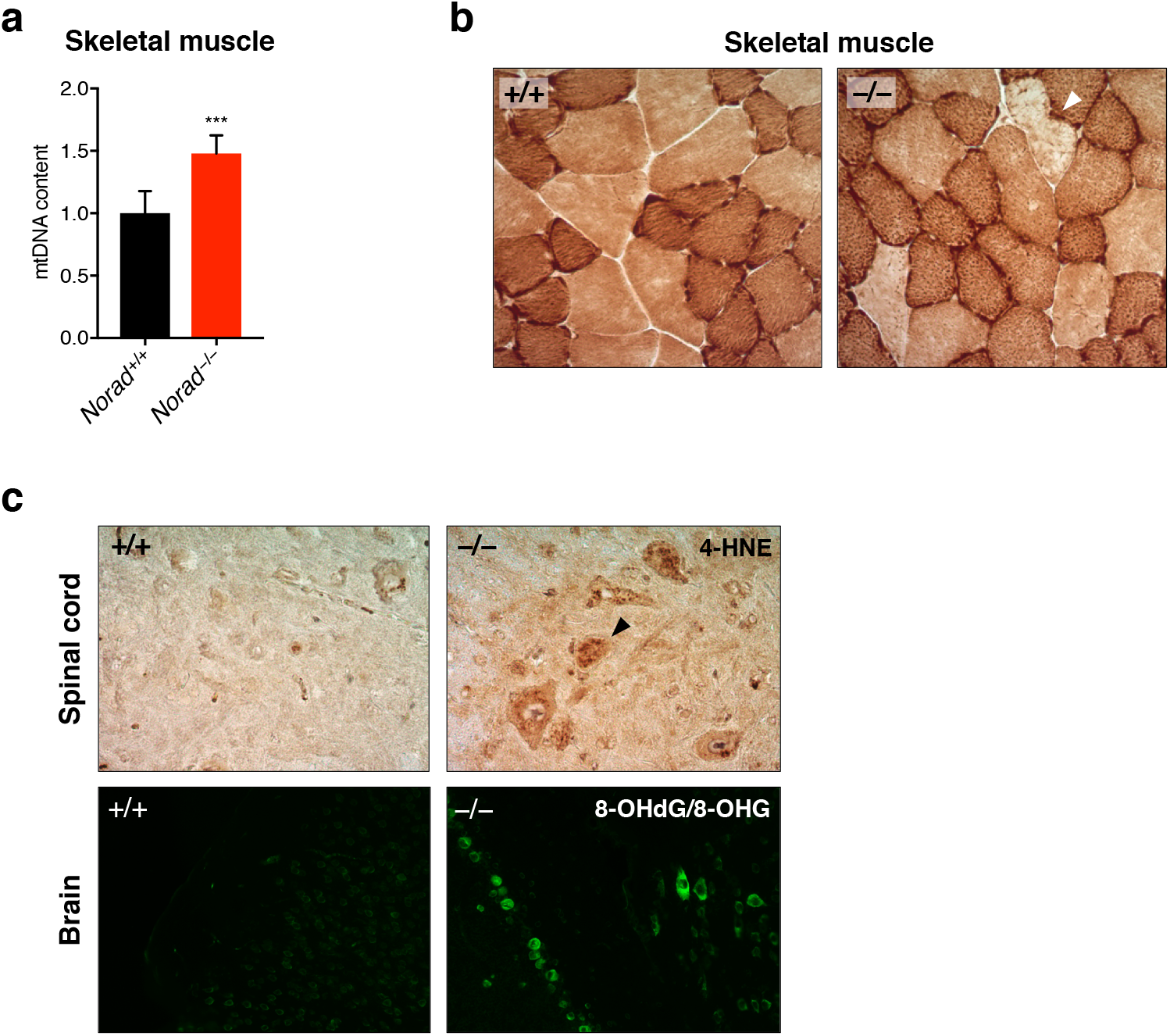
Loss of *Norad* results in mitochondrial dysfunction. **a**, Increased mtDNA content in gastrocnemius muscle of 12-month-old *Norad^-/-^* mice. mtDNA was quantified by qPCR and normalized to nuclear DNA (n = 6 mice per genotype). **b**, COX histochemistry demonstrating the presence of rare muscle fibers with greatly reduced COX activity in gastrocnemius muscle of 12-month-old *Norad^-/-^* mice (arrowhead). **c**, Oxidative damage in the CNS of 12-month-old *Norad^-/-^* mice. Lipid peroxidation and DNA/RNA oxidation were analyzed in sections of brain stem and cerebellum using 4-HNE immunohistochemistry or 8-OHdG/8-OHG immunofluorescence, respectively. Data are represented as mean ± SD in (**a**) and p value was calculated using Student’s t test. ***p ≤ 0.001.

**Supplementary Fig. 5.**
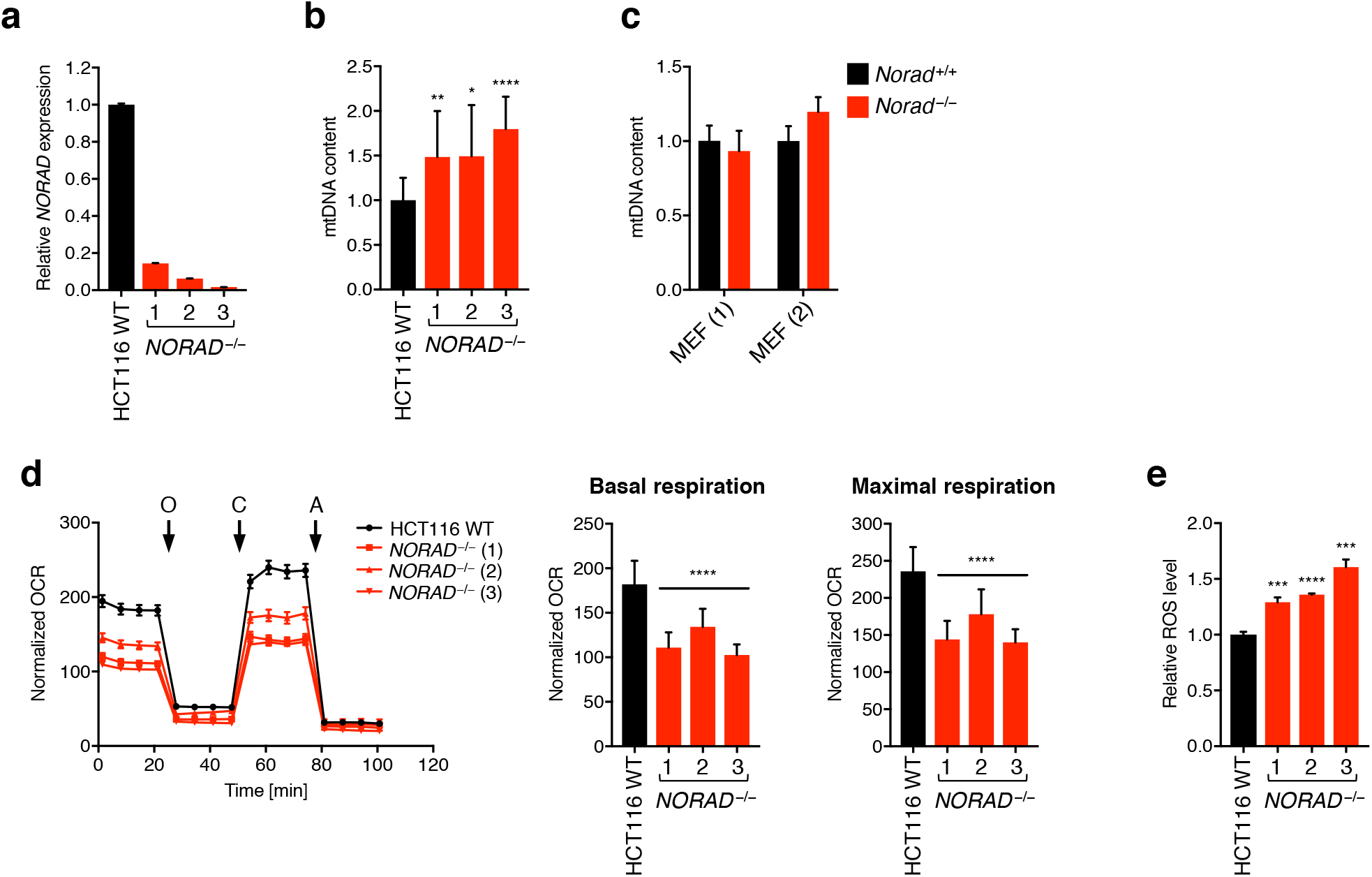
Loss of *NORAD* results in mitochondrial dysfunction and increased ROS levels in human HCT116 cells. **a**, *NORAD* levels were quantified in previously described HCT116 knockout cell lines^5^ using qRT-PCR. **b**, Increased mtDNA content in *NORAD^-/-^* HCT116 cells. mtDNA was quantified by qPCR and normalized to nuclear DNA (n = 11–13 biological replicates per condition). **c**, Immortalized *Norad^-/-^* MEFs do not exhibit elevated mtDNA content. mtDNA was quantified by qPCR and normalized to nuclear DNA (n = 3 biological replicates per MEF line). **d**, Impaired respiration in *NORAD^-/-^* HCT116 cells. Normalized oxygen consumption rate (OCR) (OCR/total protein/mtDNA content) was determined using Seahorse analysis (n = 14–16 biological replicates per condition, O = oligomycin, C = CCCP, A = antimycin A). Basal and maximal respiration was determined from measurement 4 and 12, respectively. **e**, ROS levels were assessed in the indicated cell lines using flow cytometry (n = 3 biological replicates per condition). Data are represented as mean ± SD in (**a**)-(**e**) except (**d**, left graph) which is represented as mean ± SEM for clarity due to the number of overlapping lines, and p values were calculated using Student’s t test. *p ≤ 0.05, **p ≤ 0.01, ***p ≤ 0.001, ****p ≤0.0001.

**Supplementary Fig. 6.**
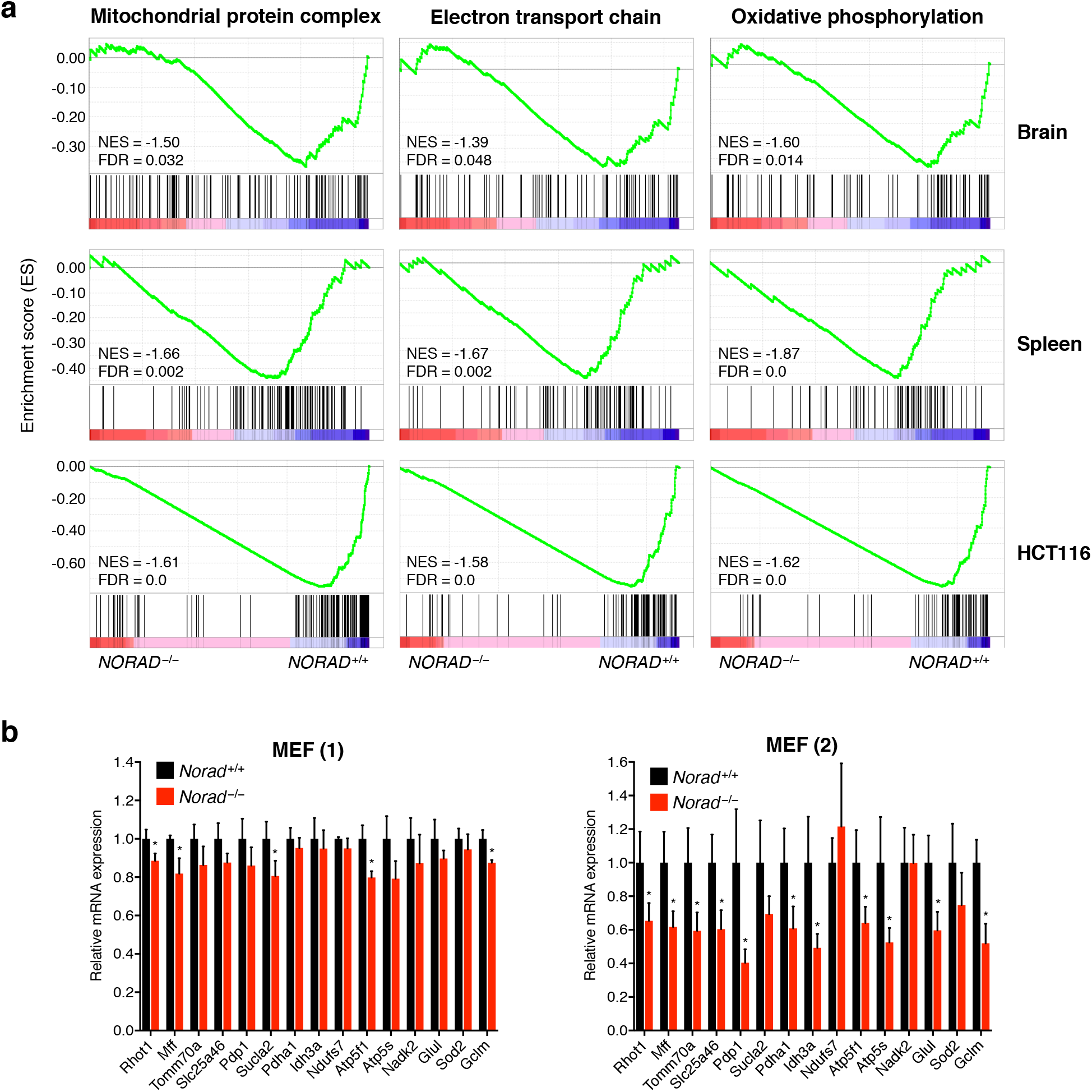
Repression of mitochondrial PUM2 target genes in *Narad^-/-^* cells and tissues. **a**, GSEA showing repression of indicated gene ontology (GO) gene sets in RNA-seq data from *Norad^-/-^* brain and spleen and human *NORAD^--^* HCT116 cells^5^. **b**, Downregulation of mitochondrial PUM2 target genes in immortalized *Norad^-/-^* MEFs. Expression levels were determined by qRT-PCR (n = 3 biological replicates for each MEF pair). Data are represented as mean ± SD in (b), and p values were calculated using Student’s t test. *p ≤ 0.05.

**Supplementary Fig. 7.**
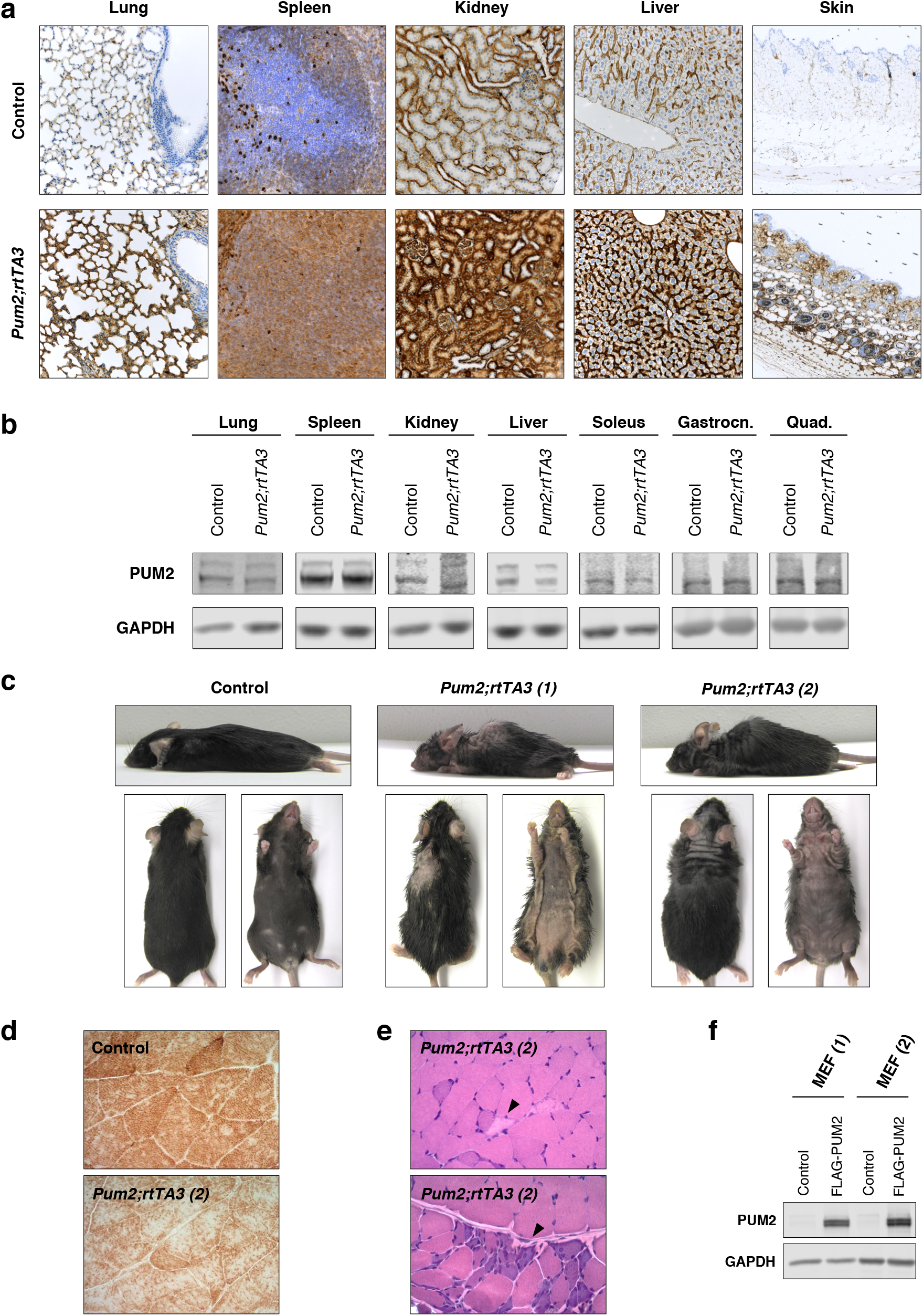
Enforced PUM2 expression phenocopies *Norad* loss of function. **a**, Images of FLAG IHC in selected tissues from *Pum2* single transgenic control or *Pum2; rtTA3* (line 1) mice after 3.5 - 6.5 weeks of dox treatment. **b**, Western blots showing total PUM2 protein levels in selected tissues from *rtTA3* single transgenic control or *Pum2; rtTA3* (line 1) mice after 3.5 weeks of dox treatment. **c**, Kyphosis and alopecia in *Pum2; rtTA3* mice from two independent transgenic founders or a wild-type control after 2 months of dox treatment. **d**, COX histochemistry showing globally reduced COX activity in gastrocnemius muscle of *Pum2; rtTA3* (line 2) mice after 2 months of dox treatment compared to a dox-treated *Pum2* single transgenic control. **e**, Histologic analysis of H&E-stained sections of gastrocnemius muscle showing necrotic fibers (upper panel, arrowhead) and basophilic regenerating fibers (lower panel, arrowhead) in dox-treated *Pum2; rtTA3* (line 2) mice. **f**, Western blot showing expression of FLAG-PUM2 in transfected immortalized wild-type MEFs.

**Supplementary Fig. 8.**
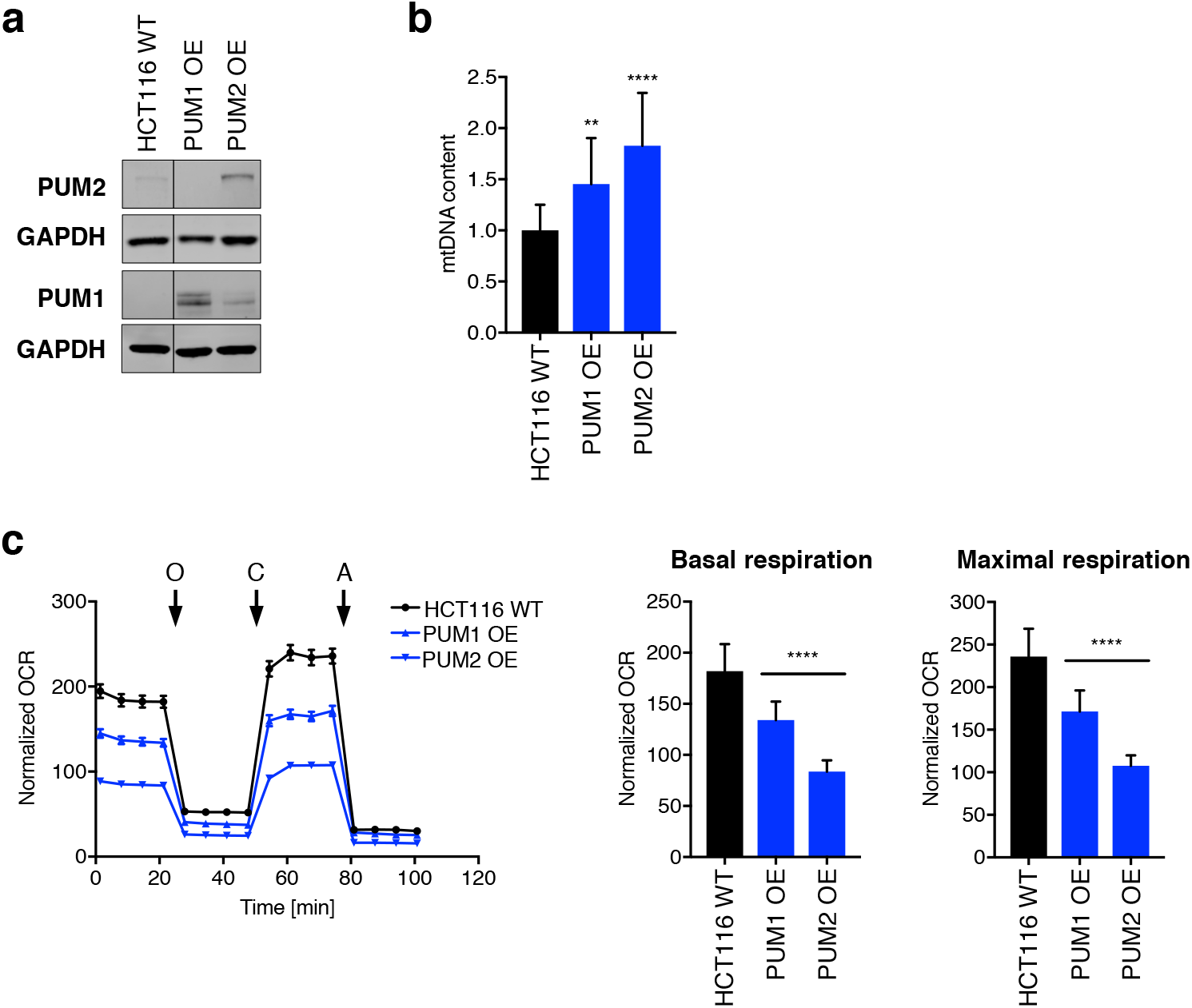
PUMILIO overexpression results in mitochondrial dysfunction in human HCT116 cells. **a**, Western blot of PUM1/2 in HCT116 cells with PUMILIO overexpression (OE). All samples originate from the same blot with irrelevant lanes cropped (indicated by the black line). **b**, Increased mtDNA content in HCT116 cells with PUM1/2 overexpression. mtDNA was quantified by qPCR and normalized to nuclear DNA (n = 11–13 biological replicates per condition). **c**, Impaired respiration in HCT116 cells with PUM1/2 overexpression. Normalized oxygen consumption rate (OCR) (OCR/total protein/mtDNA content) was determined using Seahorse analysis (n = 14–16 biological replicates per condition, O = oligomycin, C = CCCP, A = antimycin A). Basal and maximal respiration was determined from measurement 4 and 12, respectively. The quantification of the mtDNA content (b) and OCR (c) in HCT116 cells with PUM1/2 overexpression was performed at the same time as the respective experiments in *NORAD^--^* HCT116 cells (Extended Data Fig. 5b, d). Therefore, the same HCT116 WT controls as in Extended Data Fig. 5 are shown in this figure. Data are represented as mean ± SD in (b) and (c) except (c, left graph) which is represented as mean ± SEM. p values were calculated using Student’s t test. **p ≤ 0.01, ****p ≤ 0.0001.

